# Positional Interpretation of Cis-Regulatory Code and Nucleosome Organization with Deep Learning Models

**DOI:** 10.1101/2025.04.07.647613

**Authors:** Charles E. McAnany, Melanie Weilert, Grishma Mehta, Fahad Kamulegeya, Jennifer M. Gardner, Jacob Schreiber, Anshul Kundaje, Julia Zeitlinger

## Abstract

Sequence-to-function neural networks learn cis-regulatory sequence rules driving many types of genomic data. Interpreting these models to relate the sequence rules to underlying biological processes remains challenging, especially for complex genomic readouts such as MNase-seq, which maps nucleosome occupancy but is confounded by experimental bias. We introduce pairwise influence by sequence attribution (PISA), an interpretation tool that combinatorially decodes which bases contributed to the readout at a specific genomic coordinate. PISA visualizes the effects of transcription factor motifs, detects undiscovered motifs with complex contribution patterns, and reveals experimental biases. By learning the bias for MNase-seq, PISA enables unprecedented nucleosome prediction models, allowing the *de novo* discovery of nucleosome-positioning motifs and their longrange chromatin effects, as well as the design of sequences with altered nucleosome configurations. These results show that PISA is a versatile tool that expands our ability to train and interpret sequence-to-function neural networks on genomics data and understand the underlying cis-regulatory code.

## 1 Introduction

Deciphering the cis-regulatory code of gene regulation in non-coding genomic sequences is one of the remaining grand challenges in biology. A complete understanding would allow us to read the gene regulatory information in the human genome, identify genetic variants involved in disease, and design synthetic regulatory sequences for therapeutic and industrial purposes[1]. A key breakthrough to learning these cis-regulatory sequence rules are sequence-to-function neural networks[1]. These models take DNA sequence as input and are trained to predict the readout of genomics assays that measure various aspects of gene regulation, including transcription factor (TF) binding, chromatin accessibility, nucleosome organization, transcription initiation, and gene expression[2–11].

An example is the BPNet family of models, convolutional neural networks that predict the profile of genomics data at base resolution, in addition to predicting the total read counts per region[3]. BPNet was originally designed to learn high-resolution TF binding data[3, 12, 13], but the sequence-to-profile predictions and the lightweight architecture make it a robust and versatile framework for many data types, including chromatin accessibility data [14, 15] and nascent transcript data[9]. Similar architectures have successfully predicted STARR-seq/MPRA data[16] and MNase-seq nucleosome maps[17]. During training, these models learn the combinatorial interplay by which TF binding motifs generate the experimental readouts for each genomic region. For example, a model trained to predict the binding of a single TF will not only learn that TF’s binding motif, but also other sequence patterns, such as motifs for cooperative binding partners[3, 12].

Discovering novel cis-regulatory features depends, however, on effectively interpreting a trained sequenceto-function model. Neural networks have traditionally been seen as uninterpretable black boxes, but thanks to several post-hoc interpretation tools specifically designed for sequence-to-function models, sequence rules can be extracted from trained models[3, 4, 18–24]. This can be done by various attribution methods, including *in silico* saturation mutagenesis[25, 26], integrated gradients[27, 28], or corrected gradients[22]. The attribution methods deepLIFT[29] and deepSHAP[30] use Shapley values to assign each base in the input sequence a contribution score based on how much it contributed to the predicted output. Motifs are typically among the highest contributing bases due to their crucial role in the cis-regulatory code, and they are readily summarized by TF-MoDISco[31]. Rules by which motifs cooperate with each other can also be extracted from models by systematically predicting the effect of synthetic sequences *in silico*[3]. However, the motifs and their interaction rules tend to be abstract, making it challenging to deduce how motifs exert their function at individual regions. Thus, while these tools have made sequence-to-function models more interpretable, they are still limited in what they reveal.

A major drawback is that current attribution methods rely on reductive representations of how each base impacts an output prediction. Attribution methods typically quantify an input’s effects on the entire output window, and thus do not reveal where in the experimental profile a motif exerts its influence and whether the influence is narrow or broad. Furthermore, the contribution scores of motifs represent a sum of the motif’s effects and thus do not reveal whether motifs have both positive and negative effects on the predictions. For example, a motif may cause an output feature to shift to the side, thus that motif’s effect would be positive in one part and negative in another part of the experimental profile. Since mixed contributions cancel each other out, it is possible that certain motifs are missed by current attribution methods. Furthermore, some bases may be assigned contribution scores because they predict experimental biases in the data. Unless such biases are explicitly regressed out[14, 32], it is difficult to identify such contributions since the relationship to the output prediction is unclear.

Here we overcome these obstacles by introducing a new interpretation tool called pairwise influence by sequence attribution (PISA), which can be applied to sequence-to-profile models to visualize the range and level by which each individual base impacts each genomic coordinate at an individual locus. We implemented PISA in a package called BPReveal, which expands the capabilities of BPNet and ChromBPNet to support multiple different data types. We note however that PISA can be implemented in any sequence-to-profile modeling framework to visualize and interpret the learned sequence rules[2, 4, 8]. As an example, we have implemented PISA into tangermeme, a PyTorch-based version of BPNet[33].

We first describe PISA and the two types of plots it creates for individual genomic regions, squid plots and heatmaps. We then analyze previously modeled genomics data by re-training these data on BPReveal and creating PISA plots for known regulatory regions. These plots reveal details of the learned sequence rules that were previously hidden but are consistent with known biological mechanisms. This includes a motif’s influence range, previously hidden motifs that contain a mixture of positive and negative contributions, and experimental bias.

We then leverage BPReveal and PISA to train a model that predicts bias-minimized MNase-seq nucleosome data[34]. These models allow us to *de novo* discover TF motifs important for nucleosome positioning and visualize their range of effects. We further show that the effects of these motifs are bound by barriers akin to chromatin domain boundaries, which we discover *de novo* at higher resolution than Micro-C data. Finally, we show as proof-of-principle that the model can be used to generate sequence designs with altered nucleosome positioning, one of which we validate experimentally. These results pave the way to more systematically study the relationship between DNA sequence, nucleosome organization, and gene regulation.

## 2 Results

### 2.1 PISA reveals pairwise relationships between input sequence and output profile

We created PISA as an application for sequence-to-profile models, which make separate predictions for each output position (Figure 1(a)). This feature allows attribution methods such as deepSHAP[29, 30] to be applied to individual output positions, rather than for the entire predicted output window as done traditionally[3](Figure 1(b)). In our implementation, we use deepSHAP on a BPNet model and generate, for each output base *j*, contribution scores for each input base *i*. Thus, for a particular input sequence, ℙ_*ij*_ represents the Shapley value assigned to base *i* from the model’s output at base *j*. ℙ_*i*→,*j*_ represents these values in a two-dimensional matrix (Figure 1(c)).

**Fig. 1.**
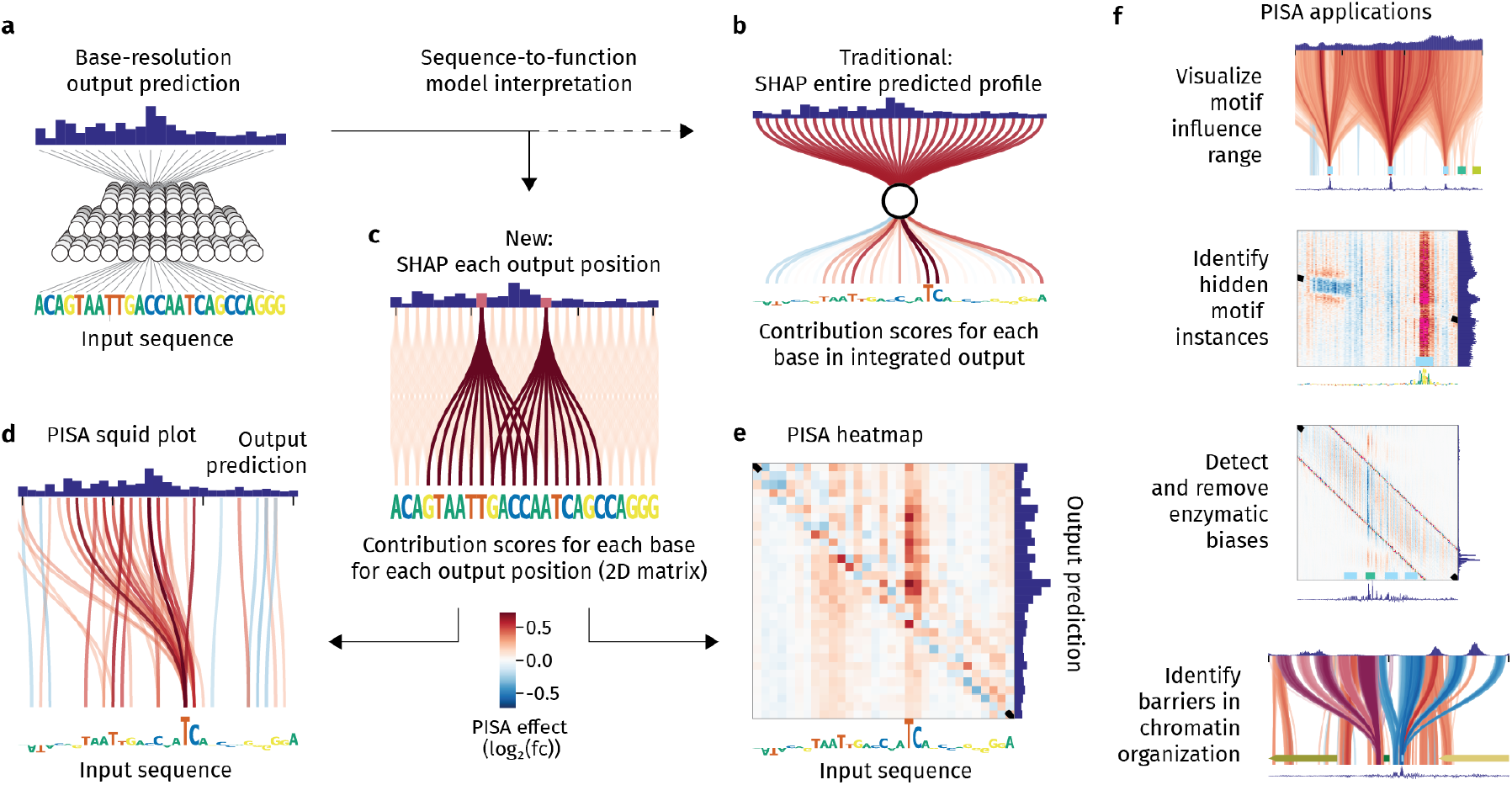
PISA and its applications. (a) A sequence-to-profile model uses DNA sequence as input and learns to predict experimental readouts from genomic assays. (b) Previous interpretation tools have assigned contribution scores to the input bases based on properties of the integrated output profile. (c) Our PISA approach assigns contribution scores for each output individually, resulting in a 2D matrix ℙ_*i*→*j*_ . (d) Squid plots show high PISA values as colored lines that connect an input base to an output position. In this example, the TCA motif has a strong effect, and the bulk of this effect is to the left of the motif’s position. (e) A heatmap of the same region used in (d) shows PISA values as a colored grid. While less immediately readable than the squid plots, heatmaps are useful when multiple effects overlap or when experimental biases are present at the diagonal. (f) PISA is implemented in the BPReveal package and has multiple applications further described in this study.

To visualize this matrix, we made two types of PISA plots. In the **PISA squid plot** (Figure 1(d)), we sought to create a simple summary of the range by which sequence patterns impact the output signal. The input bases are displayed at the bottom of the squid plot as traditional contribution scores, with lines drawn from base *i* at the bottom to base *j* in the predicted output profile at the top. The color of the line represents the contribution score ℙ_*i* → *j*_ (positive in red, negative in blue). PISA values below a threshold are not shown for clarity.

The **PISA heatmap** (Figure 1(e)) provides a finer-grained representation, where the entire PISA matrix is visualized as a colored grid: The pixel at position *i* on the x-axis and position *j* on the y-axis represents ℙ_*i* → *j*_ (positive in red, negative in blue). For reference, the traditional contribution scores are again shown at the bottom, while the output prediction is shown on the right. In this grid, the diagonal is where the input base affects the output at the same position (i.e., *i* ≈ *j*). As we will see later, this diagonal of local influence is often where experimental bias is found.

To leverage PISA and its downstream applications (Figure 1(f)), we implemented a package called BPReveal. Since it encompasses the capabilities of BPNet[3] and ChromBPNet[14], BPReveal models can make use of any combination of multi-head, multi-strand architectures and leverage separate bias models to remove experimental biases from data (see architecture in Extended Data Figure 1).

### 2.2 PISA visualizes the influence range of TF motifs

To visualize how TF motifs influence output predictions, we first applied PISA to TF binding data in an already-characterized system (Figure 2(a,b)). We trained a BPReveal model on previously published high-resolution ChIP-nexus data of Oct4, Sox2, Klf4, and Nanog in mouse embryonic stem cells[3], which gave results on par with the original BPNet model (Extended Data Table 1). We then generated PISA squid plots for Oct4 and Nanog binding at a key pluripotency enhancer, *Lefty1*, which revealed distinct influence ranges and a hierarchical relationship between the *Oct4-Sox2* motif and the three *Nanog* motifs (Figure 2(a,b)).

**Fig. 2.**
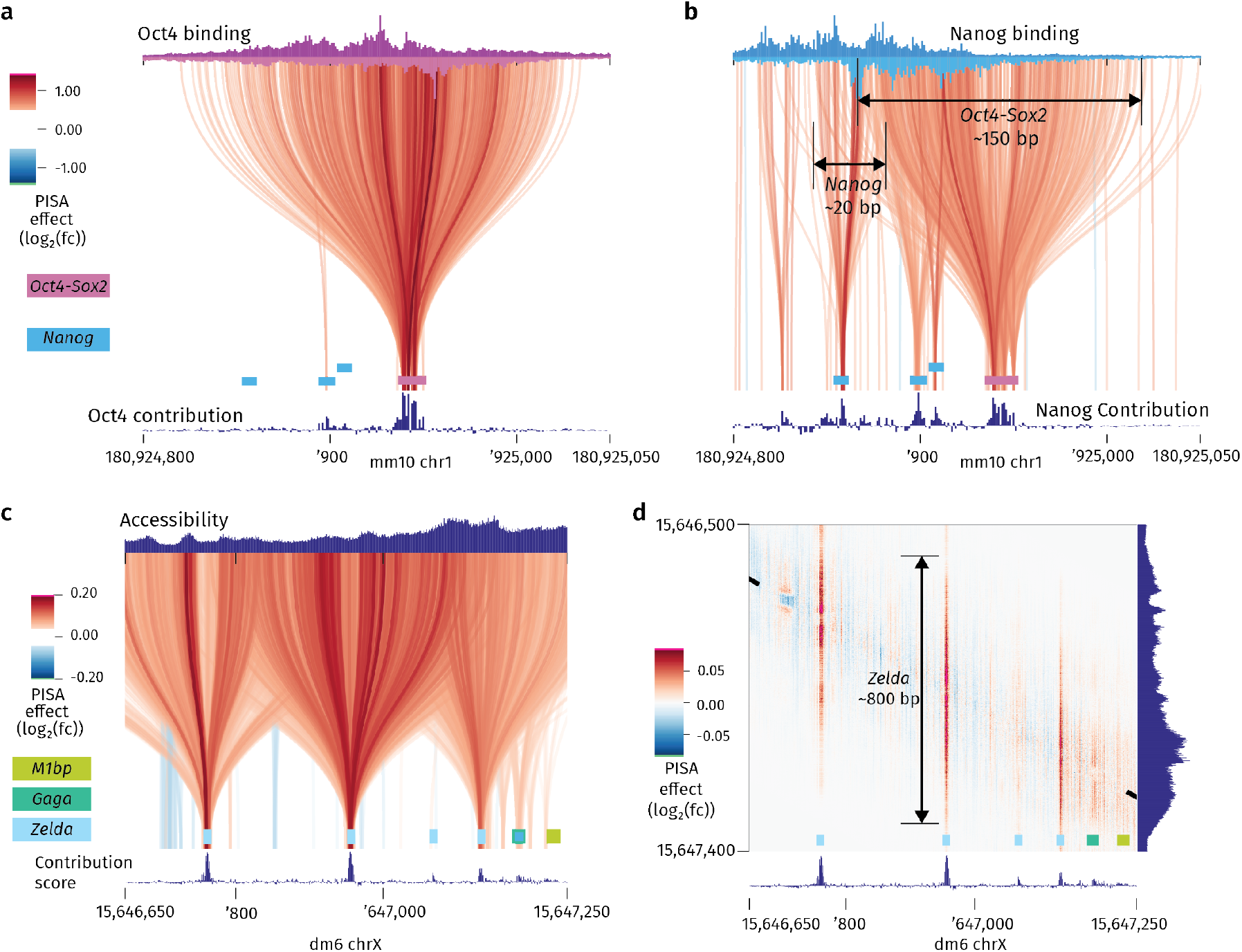
Visualizing the influence range of motifs using PISA. (a) Squid plot showing that the *Oct4-Sox2* motif (purple) determines Oct4 binding in a broad window. A multi-head model was trained on stranded high-resolution binding data (ChIP-nexus) for the pluripotency TFs Oct4, Sox2, Klf4, and Nanog. The predicted Oct4 ChIP-nexus data at the *Lefty1* enhancer are shown at the top (plus strand in dark purple, minus strand in light purple). The PISA squid plots (both strands are overlaid) show that the Oct4 task assigns broad importance to the *Oct4-Sox2* motif, consistent with Oct4 being a pioneer TF whose activity is not governed by other nearby motifs. (b) Squid plot of the same region as (a), showing that *Nanog* motifs (blue) promote Nanog binding in a narrow range, while the *Oct4-Sox2* motif (pink) has a broad influence. To demonstrate the influence of the *Oct4-Sox2* motif, a separate model was trained only on Nanog binding. The predicted Nanog ChIP-nexus data at the *Lefty1* enhancer are shown at the top (plus strand in dark blue, minus strand in light blue). The broad effect of the *Oct4-Sox2* motif and the narrow effect of the *Nanog* motifs are consistent with Oct4 and Sox2 being pioneering TFs, while Nanog is not. (c) Squid plot of the *sog* enhancer from a model trained on bias-minimized ATAC-seq data from fly embryos[15]. The chromatin accessibility is largely determined by three motifs for the pioneer factor Zelda (turquoise). While aesthetically pleasing, the squid plot is difficult to interpret due to the many overlapping lines. (d) A PISA heatmap shows the overlapping effects of the three *Zelda* motifs more clearly, and reveals that the central motif drives chromatin accessibility over a window of ∼800 bp.

The PISA squid plot for Oct4 binding (Figure 2(a)) showed that the *Oct4-Sox2* motif directs the Oct4 binding footprint in a broad window of ∼150 bp. The *Nanog* motifs show no contribution to Oct4 binding. The PISA squid plot for Nanog binding (Figure 2(b)) shows that the three *Nanog* motifs contribute to Nanog binding more locally, while the *Oct4-Sox2* motif promotes Nanog binding in a broader window. The broad effect of *Oct4-Sox2* and the local effect of *Nanog* are likely because Oct4 and Sox2, but not Nanog, are pioneer TFs in mouse embryonic stem cells[35, 36].

To illustrate the applicability of PISA in a different data type, we next trained a BPReveal model to predict bias-minimized ATAC-seq chromatin accessibility data from *Drosophila*[15] (Figure 2(c,d)). Our model yields results on par with the previously published ChromBPNet model (Extended Data Table 2) and rediscovered the *Zelda* motif.

Using the well-studied *sog* shadow enhancer as example, the PISA squid plot showed each *Zelda* motif as a wide squid-like pattern reaching a region of several hundreds of bases around the motif (Figure 2(c)). Likewise, the heatmap shows an influence range of ∼800 bp for the central *Zelda* motif (Figure 2(d)), consistent with Zelda being a key pioneer TF in the early *Drosophila* embryo[15, 37, 38].

We conclude that PISA directly visualizes the influence range of motifs at single loci, providing clues to the effect of those motifs on chromatin state.

### 2.3 PISA reveals complex motif effects

We next used PISA to explore motifs that influence the output signal positively at some parts of the region, while having a negative influence in other parts. When using traditional attribution methods, such mixed effects might cancel each other out, causing these motifs to be misrepresented or difficult to discover.

Serendipitously, the PISA heatmap of the chromatin accessibility at the *sog* shadow enhancer provided such a pattern (left edge of Figure 2(d), enlarged in Figure 3(a, left)). We observed a pattern with central negative contributions (blue) flanked by positive contributions (red), mirroring the predicted output profile at this position, i.e. a pronounced central dip flanked by peaks on each side, as shown on the right (Figure 3(a, left)). The mixed negative and positive contributions nullify each other since the counts contribution track below shows no contribution (shown below with an arrow).

**Fig. 3.**
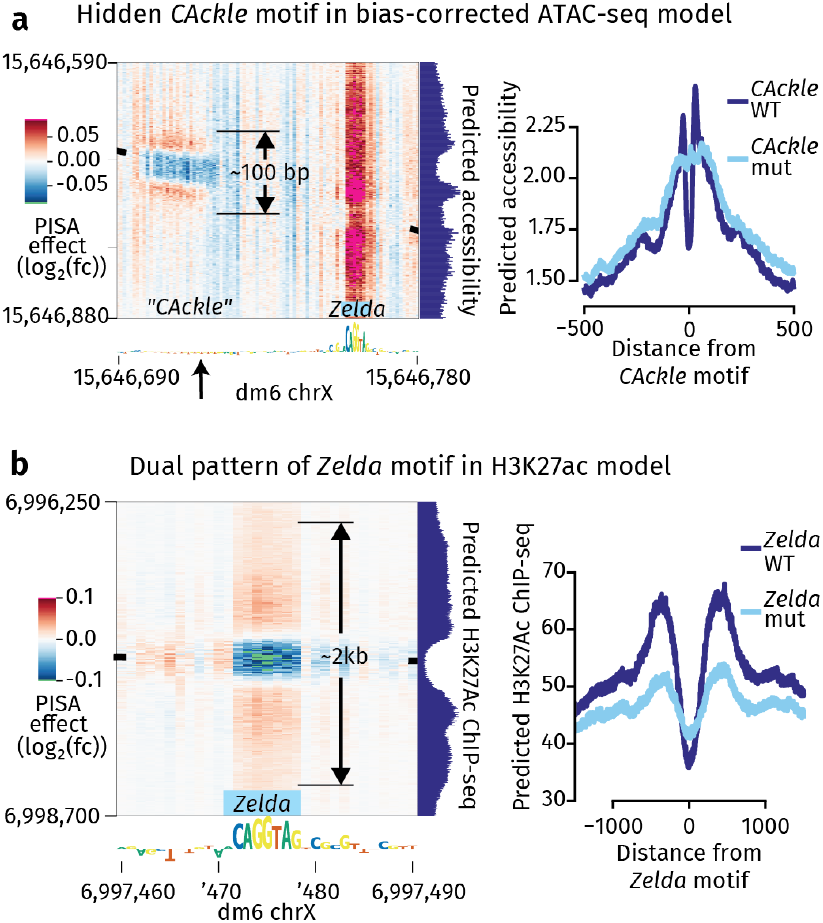
Two motifs with positive and negative effects. (a) A CA repeat creates a local dip in chromatin accessibility in ATAC-seq. (Left) The CA repeat motif, which we refer to as *CAckle*, has both negative (blue) and positive (red) contributions in a PISA heatmap, and thus is not detected in the counts contribution score (arrow). It creates a small (∼50 bp) dip in accessibility surrounded by a shoulder of slightly higher accessibility, as seen in the predicted output profile on the right. The motif’s entire effect range is on the order of 100 bp, which is much smaller than the ∼800 bp effect of the *Zelda* motif on the right of the heatmap. (Right) We simulated the effects of mutating *CAckle* motifs, and found that replacing the CA repeat with random nucleotides abolished the local dip and shoulder effect, but had a minimal effect on the global counts. (b) Positive and negative effects are a signature of histone modifications. (Left) A model trained on ChIP-seq data for H3K27Ac, a marker of enhancer activation, shows a similar negative-and-positive pattern around a *Zelda* motif. This is consistent with Zelda causing the nucleosomes in an enhancer to be depleted (hence the central dip) and neighboring nucleosomes to be acetylated (hence the positive shoulder). Unlike the short-range *CAckle* effect, the *Zelda* motif drives acetylation in a 2 kb window. (Right) Perturbing many instances of the *Zelda* motif shows the same effect genome-wide: Mutating the *Zelda* motif decreases both the depth of the central dip and the acetylation of flanking nucleosomes.

The sequences that produce this pattern correspond to a `CA` repeat (referred to as *CAckle*), which has previously been shown to boost TF binding and enhancer activity[13, 39]. Although *CAckle* sites have dual positive and negative contributions, the total effect is in some genomic instances not neutral, allowing us to discover them across the genome using TF-MoDISco and motif scanning. To test their effect, we performed simulated mutations of 712 mapped *CAckle* motifs (Figure 3(a, right)). Upon mutation, the dip with the flanking peaks flattened out, but the overall read counts remained similar, confirming a mixed negative and positive effect of *CAckle* motifs on the chromatin accessibility. Although a *CAckle*-like motif also contributes to the Tn5 enzymatic bias, the central depletion around instances of the *CAckle* motif is also reflected in experimental ATAC-seq data (Extended Data Figure 2).

Another example of a complex motif effect is provided by ChIP-seq data of H3K27ac, a histone modification that flanks active enhancers. We trained a model on H3K27ac ChIP-seq data from early *Drosophila* embryos[15] and visualized known enhancers using PISA. This revealed that the H3K27ac profile predictions strongly depend on *Zelda* motifs (Figure 3(b, left)). A *Zelda* motif creates a dip in H3K27ac at the center of the enhancer, seen as negative contributions (blue), while also promoting H3K27ac at the enhancer’s flanks, seen as positive contributions (red). This trend was again confirmed by mutating 325 *Zelda* motifs, which decreased H3K27ac at the flanks and made the central dip more shallow (Figure 3(b, right)). This dual effect is consistent with Zelda’s role as pioneer TF that depletes nucleosomes in the center, while recruiting the acetyltransferase Nejire to acetylate the flanking nucleosomes[40, 41].

The dual pattern of the *Zelda* motif in the H3K27ac model is similar to that of the *CAckle* motif in the chromatin accessibility model, but the range of the effect is an order of magnitude larger. While the contributions of *CAckle* spanned ∼100 bp centered on its motif, those of *Zelda* extended to a ∼2 kb region. These results show how PISA provides additional spatial resolution for interpreting how motifs exert their effect.

### 2.4 PISA detects experimental biases and enables bias-corrected MNase-seq models

If PISA is able to identify how each base positionally impacts the output predictions, it should visualize experimental biases, which tend to be local (Figure 4). For example, ATAC-seq uses the transposase Tn5, which has a local preference towards certain motif-like sequences[42]. This problem has been elegantly solved in a package called ChromBPNet, where a separate BPNet bias model is trained on data of closed regions that contain the bias but minimal accessibility[14]. The frozen bias model is then used to train a new BPNet model which learns the residual signal that must be added to the bias in order to predict the experimental profile. In this way, the experimental bias and regulatory sequence rules are separated, partitioned into distinct models for downstream interpretation.

**Fig. 4.**
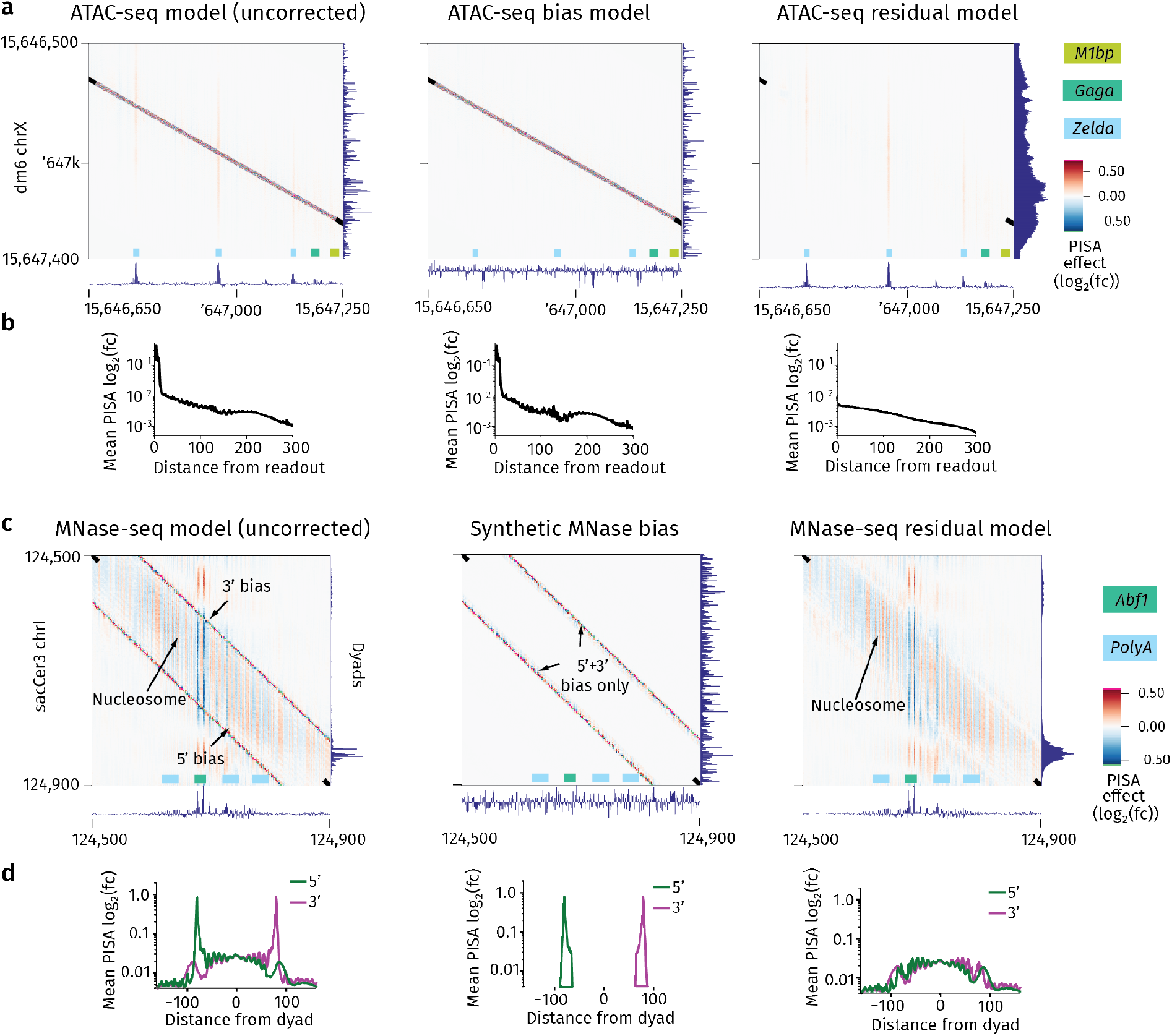
PISA reveals enzymatic biases. (a) The ChromBPNet bias correction strategy effectively removes experimental bias. (Left) A model trained on ATAC-seq endpoints in accessible regions learns the effects of motifs (faint vertical bands), but also learns enzymatic bias (strong diagonal band). (Center) A model trained on closed regions of chromatin learns enzymatic bias, but not pioneering motifs. (Right) The ChromBPNet architecture removes the patterns learned by the bias-only model and assigns importance only to the pioneering motifs. The tracks below all PISA heatmaps show the read count contributions, which captures the total accessible signal well, while the profile contributions more strongly capture the experimental bias (Extended Data Figure 3). (b) A base attribution total (BAT) plot quantifies the strength of the bias. The first two models show a large spike in contribution near the diagonal of the PISA heatmap, while the right plot shows no such central spike. (c) PISA enables bias-minimized MNase-seq models. (left) A model trained on MNase endpoints learns the enzymatic bias twice: Once on the 5^*′*^ end of the nucleosome-sized fragment and once again on the 3^*′*^ end. By aligning and subtracting the PISA values for each strand (see section 4.5), we construct a synthetic bias track that can be used to correct MNase sequence bias using the ChromBPNet architecture. Tracks below all PISA heatmaps show the tracks for profile contributions, not count contributions, since the total read counts do not substantially change across regions when predicting nucleosome profiles. (d) A BAT plot quantifies the effectiveness of MNase bias removal. The large spikes of importance at +80 and -80 are enzymatic bias, and these spikes are eliminated in the bias-corrected model (right).

To visualize the Tn5 bias, we inspected the BPReveal models we trained to predict the bias-minimized ATAC-seq data from the early *Drosophila* embryo[15], using the *sog* shadow enhancer as an example (Figure 4(a)). In PISA heatmaps of the combined model that predicts the uncorrected data, the ATAC-seq bias is visible as a diagonal band that overshadows the weaker effects of the three *Zelda* motifs that appear as vertical stripes (Figure 4(a, left)). In the bias model, all that is left is the diagonal band (Figure 4(a, center)), while the residual (i.e., bias-minimized) model only shows the effects of the three *Zelda* motifs (Figure 4(a, right)). This confirms that the diagonal band is indeed the bias and that the bias was successfully removed in the residual model.

As a method for quantifying the effectiveness of the bias removal, we created a base attribution total (BAT) plot, which shows the average contribution of each input base to the output at a given distance (Figure 4(b)). More precisely, BAT(Δ) = mean_*i*_(ℙ_*i*→ *i*+Δ_), where Δ is the x-axis in the heatmap. For the bias model and the uncorrected ATAC-seq model, the BAT plot shows the local bias as 100-fold increase within 10 bp (Figure 4(b left, center)), while this signal is not present in the bias-corrected model (Figure 4(b, right)).

Having confirmed that PISA reveals experimental bias in BPReveal model predictions, we next examined other data types. Since some data types lack appropriate control regions for training a bias model, PISA may even help create an experimental bias track. A good example are nucleosome maps created by MNase-seq, which have a strong AT sequence bias[34, 43]. However, nucleosomes are present across the genome and so there are no regions that could be used to train a bias model[44].

We trained a BPReveal model on MNase-seq data from *S. cerevisiae* since yeast nucleosome occupancy is well-studied[45–47], the small genome size permits very high coverage data[48], and we can benchmark our results against previous deep learning models[25]. Since the MNase enzymatic bias is found at the fragments’ end, we recorded each fragment’s 5^*′*^ and 3^*′*^ ends as two separate tracks and trained BPReveal to predict both tracks simultaneously (see section 4.5).

BPReveal achieved high prediction accuracy on experimental data from held-out chromosomes (Extended Data Table 3). It achieved higher prediction accuracies than the model by Routhier et al[25], using the same training data. This was true even when the model performance was scored by Pearson correlations, which are part of the loss function that the Routhier model optimizes during training (Extended Data Table 4). This suggests that BPNet-derived models trained with the BPReveal framework are well-suited to learning MNase-seq data in yeast.

We then generated PISA heatmaps to assess whether we could identify the experimental bias of MNase-seq. PISA heatmaps showed a strong, narrow diagonal band, either to the left (in the 5^*′*^ model) or the right (in the 3^*′*^ model) of the nucleosome-sized fragments (Figure 4(c, left)). This signal represents highly local effects in BAT plots (Figure 4(d, left)) and thus, as with ATAC-seq data, local enzymatic sequence bias strongly contributes to the predictions. Apart from the bias, the PISA heatmaps also showed subtle positive and negative contribution scores within the nucleosome-sized fragments (shown combined in Figure 4(c, left)). This signal likely represents intrinsic DNA sequence properties that determine the readiness to wrap around the histone core and form stable nucleosomes[47, 49–52].

Reasoning that a bias-minimized MNase-seq model would predict a cleaner nucleosome profile and enhance interpretation[14], we explored ways to extract the bias from the PISA values and derive a synthetic bias track. The model’s independent prediction of the 5^*′*^ and 3^*′*^ ends serendipitously provided us with a method to extract the bias. Since the PISA heatmaps of the 5^*′*^ and 3^*′*^ ends differ in the bias but essentially not in the biological signal, subtracting the two maps cancels out the biological portions and leaves behind pure enzymatic bias. The effectiveness of this approach is seen in BAT plots (Figure 4(d)).

The second innovation came from the insight that the efficiency property of Shapley values allows us to sum up the PISA values from the bias and construct a genome-wide synthetic MNase bias track (Section 4.5). With this track, we then trained an MNase bias model and confirmed (using TF-MoDISco) that we did not learn any TF motif (Extended Data Figure 4).

This bias model then allowed us to train and cross-validate a combined model that captures the bias-minimized nucleosome signal in the residual model. PISA heatmaps confirmed that we have successfully separated the MNase-seq bias and nucleosome signal (Figure 4(c)).

To benchmark the bias removal, we compared our results to those of other MNase bias removal methods. We trained a bias model on MNase-seq data obtained from naked DNA[48, 53] or used statistical modeling such as seqOutBias[32] before training on the MNase-seq data. Both of these methods left more uncorrected enzymatic bias than our corrected prediction (Extended Data Figure 5). Thus, PISA not only visualized the bias in MNase-seq data, but allowed us to derive a synthetic bias track to train a bias-minimized MNase-seq model.

### 2.5 PISA visualizes how *de novo* discovered motifs affect nucleosomes

We next examined what the MNase-seq model learned. For an intuitive reading of nucleosome centers (dyads), we displayed the MNase-seq data as midpoints of the nucleosome-sized fragments, rather than the positions of the 5^*′*^ and 3^*′*^ ends that we trained the model on (Section 4.1.3). As seen at the *Ade1* gene example locus, a known genetic marker in yeast, the experimental and predicted midpoint data look spiky. Only after removal of the bias, the nucleosome signal was notably smoother, and the background signal in the profile contribution scores was markedly reduced, revealing motifs such as *Reb1* (Figure 5(a)).

**Fig. 5.**
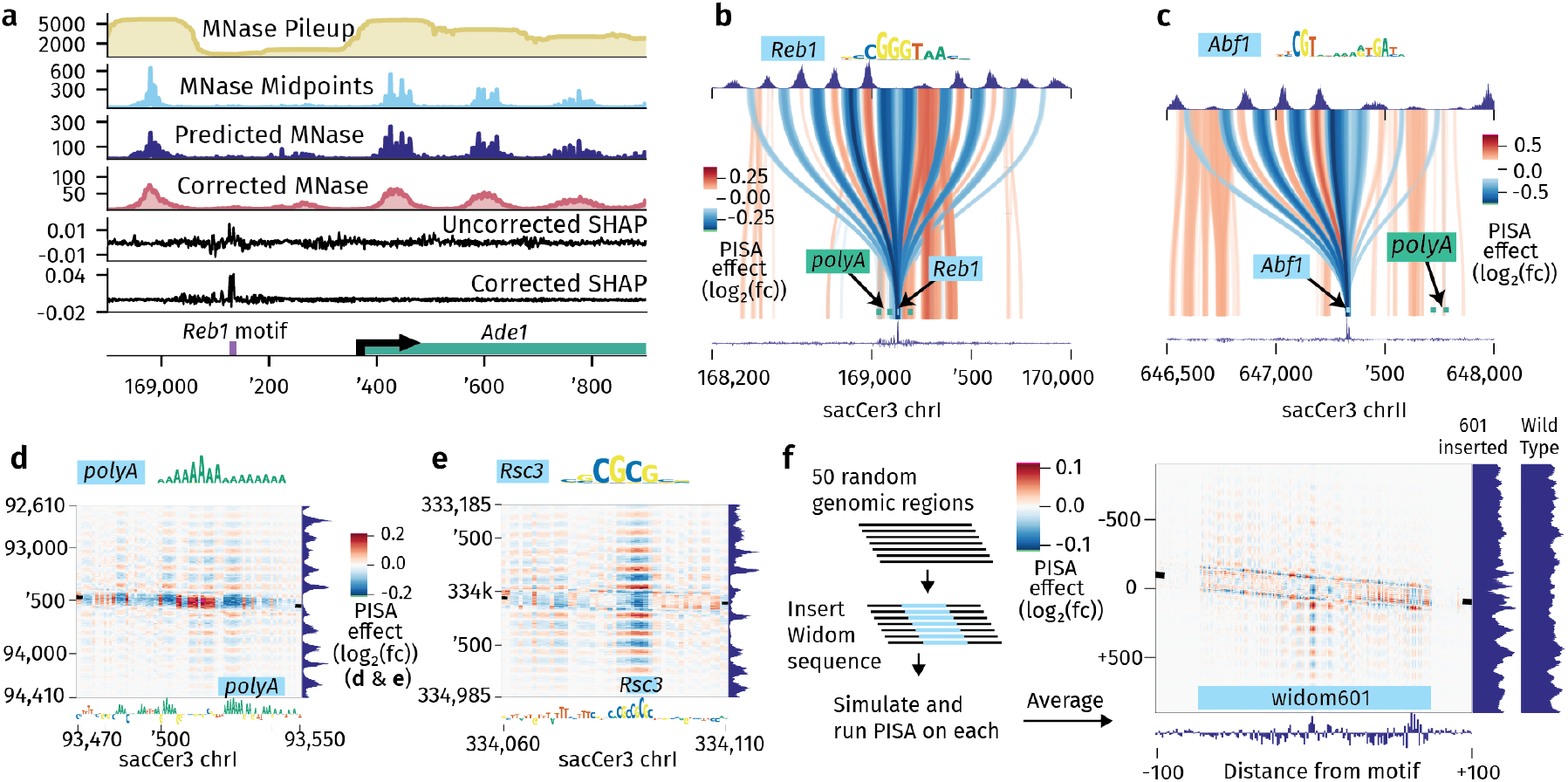
The bias-corrected MNase-seq model discovers motifs *de novo* that positions nucleosomes. (a) The model learns and corrects MNase data. The top track shows the MNase-seq data as fragment pileup, a common way of plotting such data. Our models instead predict the endpoints of the fragments, providing higher resolution data. For convenience, we present the model’s predictions as midpoints. The model’s predictions match the experimental data well, and the corrected data remove much of the high-frequency noise in the uncorrected data. The importance of this correction is most visible in the contribution scores, which are noisy and fuzzy in the uncorrected model but clearly show a *Reb1* motif in the corrected model. (b) A PISA squid plot of the region in (a). The central *Reb1* motif shows a long-range ringing effect. (c) A representative *Abf1* motif shows a similar long-range ringing effect. This particular motif instance shows a striking leaning effect, where the motif drives nucleosome positioning mostly to its left. (d) The *polyA* motif is more subtle than *Reb1* or *Abf1*. It tends to be more distributed, as in this example where a block of *polyA* has been annotated by TF-MoDISco, but several smaller clusters of A also contribute to position nearby nucleosomes. (e)The *Rsc3* motif also creates a ringing effect. In this instance, it is coupled with several short T repeats, which likely serve as a very degenerate *polyA*. (f) The widelyused Widom 601 sequence that strongly positions nucleosomes *in vitro* does not have a strong effect *in vivo*, consistent with previous observation[54]. PISA was applied to 50 random genomic regions where the Widom 601 sequence was inserted and then averaged. The average output prediction (601 inserted) is also shown to the right, in comparison to those of the wild-type sequences. The effect is, on the whole, very small. The complete analysis showing all plot types for all motifs is in Extended Data Figure 7.

We then ran TF-MoDISco to discover motifs *de novo* using either the contribution scores from the uncorrected model, the bias-only model, or the bias-corrected model. The bias-only model learned many motifs related to the MNase sequence bias, most notably sequences that transition from AT-rich to GC-rich regions, consistent with previous studies[34, 43] (Extended Data Figure 4). The uncorrected model also learned biologically relevant motifs, but it was difficult to distinguish them from the many bias motifs it had also learned. In contrast, the motifs discovered from the bias-corrected model were recognizably biologically relevant, with motifs known to affect nucleosomes such as *Abf1* and *Reb1, polyA* repeats, CG-rich sequences, and a CGCG motif[46, 55–57].

While these motifs were known, no neural network model has previously *de novo* discovered and mapped these motifs genome-wide, or helped provide insights into their effect sizes and ranges [17, 25, 58]. The CGCG motif has been characterized as a recognition signal for the RSC remodeler complex[59, 60], but the frequency of such motifs in the genome is unknown. *polyA* sequences resist DNA bending and are often found between nucleosomes[61–63], but the exact sequence nature and context by which such sequences affect nucleosomes is not clear. We therefore scanned the genome for motifs contributing to the nucleosome predictions and visualized their effects with PISA. As a control, we compared our discovered motifs to those discovered by traditional motif scanning (methods).

PISA squid plots showed a striking effect of *Reb1* and *Abf1* motifs (Figure 5(b,c)). Both induce a strong oscillating positional signal of ∼750 bp (or 4-5 nucleosomes) emanating from the motif, usually extending in both directions but sometimes leaning towards one side. We discovered 656 *Abf1* motifs and 760 *Reb1* motifs, the majority of which were discoverable with traditional motif scanning (Extended Data Figure 6). By contrast, the 1,543 *Rsc3* (CGCG) and 12,791 *polyA* motifs mapped by the model could not be identified by traditional motif scanning without discovering a large number of false positive motifs not associated with nucleosome positioning (Extended Data Figure 6). This suggests that the model learned finer-grained sequence context not represented in the motifs. Consistent with this, PISA squid plots and PISA heatmaps showed that the positioning effects of *Rsc3* motifs, and especially those of *polyA* motifs, were weaker, less localized and more enigmatic than those of *Reb1* and *Abf1* motifs (Figure 5(d), Extended Data Figure 7). Since the weak CGCG and *polyA* motifs were very abundant across the genome, their combined effect nevertheless made a substantial contribution to the nucleosome signal (Extended Data Figure 6).

To put these effects into context, we asked what pattern the model would predict for the Widom 601 sequence, a ∼150bp-long sequence often used by *in vitro* experiments to position nucleosomes[64]. To provide a neutral sequence context, we injected the 601 sequence into 50 random sequences, made predictions, performed PISA, and averaged the results (Figure 5(f)). The Widom 601 sequence showed even smaller oscillating long-range effects than those of *polyA* motifs and showed predominantly intrinsic (short-range) effects. This is consistent with previous experimental evidence suggesting that the Widom 601 sequence does not position nucleosomes *in vivo*[54] and points to differences between *in vitro* and *in vivo* experiments.

In summary, our model interpretation revealed that motifs play a major role in positioning nucleosomes across the genome, which is surprising given that previous debates in the field have mostly focused on the intrinsic effects of DNA sequence[47, 55, 56, 65–70].

### 2.6 Barrier elements limit the effects of nucleosome-positioning motifs

PISA plots showed that a motif’s long-range nucleosome positioning effect, particularly around promoters, often leans to the left or right (Figure 5(c)). We hypothesized that our model has learned barrier elements of chromatin organization. Given the close link between nucleosome positioning and 3D chromatin interactions in yeast[72], these barriers could correspond to the domain boundaries seen in Micro-C data[71]. If domain boundaries block the effect of nucleosome-positioning motifs, we would expect to identify them if we systematically insert nucleosome-positioning motifs at various positions along a genomic sequence (Figure 6(a)). When the effect of the motif is attenuated on one side, we quantify the “lean effect” as the difference between the effect on the left versus the right (Figure 6(a)).

**Fig. 6.**
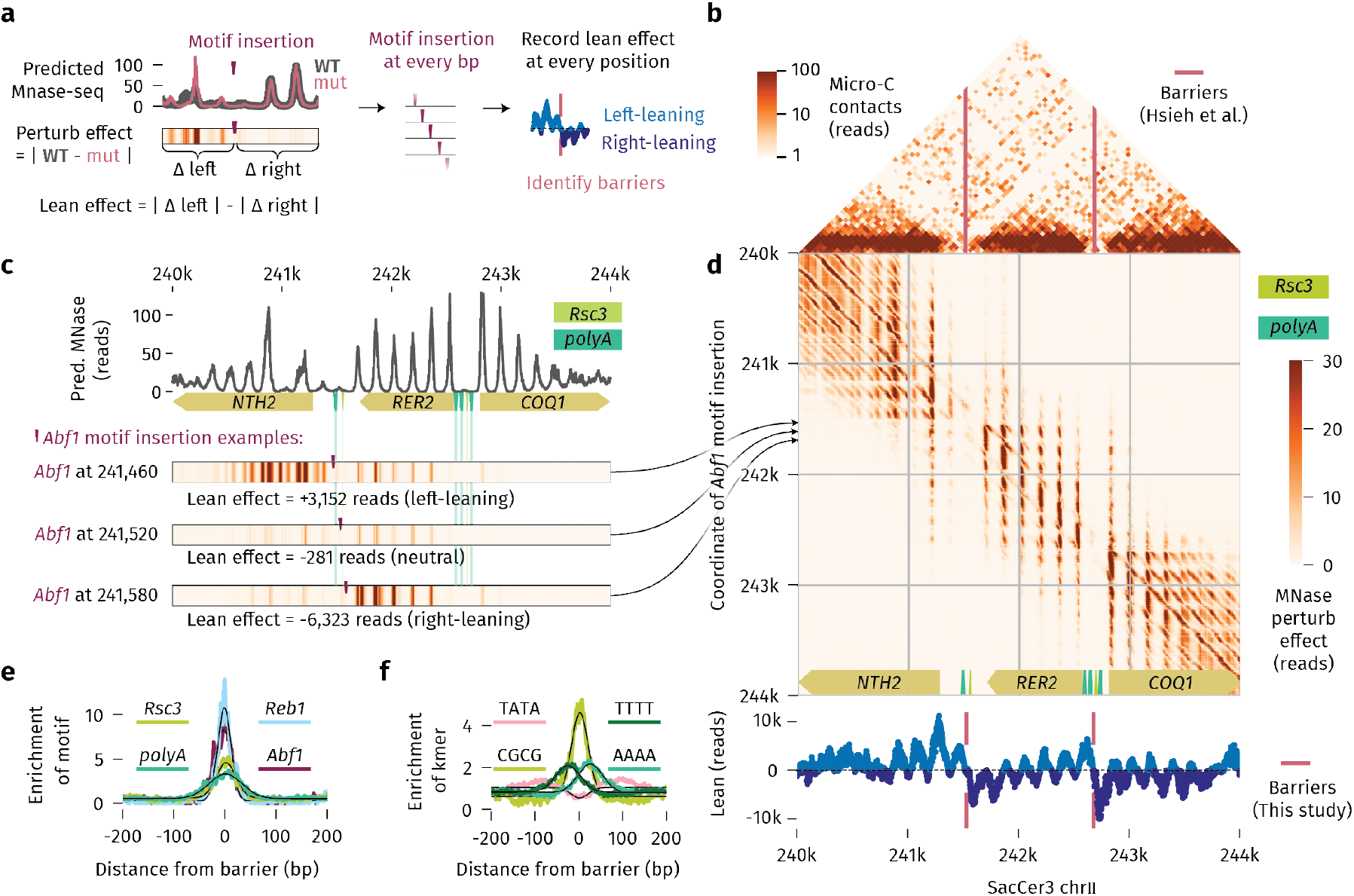
A model for nucleosome positioning learns chromatin organization. (a) Process for identifying domain boundary elements. (left) After inserting a strong positioning motif (purple pip), the changes in predicted nucleosome positions (pink line) from the wild-type prediction (gray) are recorded. The difference is summarized below as a heatstrip, showing that most effects are left-leaning, resulting in a positive lean value. (center) The motif is injected at every possible position in the genome to record the lean effects. (right) Positions where lean values flip from positive (cyan) to negative (navy) represent a barrier (pink). (b) Published Micro-C data[71] with identified barriers (pink) for a region around the *RER2* gene, which is part of the test set in our MNase-seq model. (c) Example insertions from the *RER2* region. (top) The wild-type MNase-seq prediction shows a wide nucleosome-depleted region between *NTH2* and *RER2*, a promoter region that contains mapped motifs. (bottom) Insertion of an *Abf1* motif at three positions (purple ticks) leads to dramatically different effects as seen on the heatstrips. The effect is left-leaning at position 241,460, overall weak at position 241,520, and right-leaning at position 241,580. This suggests that sequence elements near 241,520 limit the effect of motif insertions. (d) (top) When the motif is systematically injected at every position in the *RER2* region, the resulting heat map resembles the Micro-C contacts shown in (b). (bottom) Lean effects for every motif insertion show positive values (cyan) for left-leaning and negative values (navy) for right-leaning effects. At a barrier (pink), the lean values flip from positive to negative. (e) Motif enrichments at all identified barriers in the yeast genome show that the *Abf1* and *Reb1* motifs are strongly enriched within 20 bp of the barrier positions. The *polyA* motif is similarly enriched, but over a broader window of 30 bp, likely because *polyA* has an asymmetric effect and motif direction was not considered in this plot. (f) Short sequence elements enriched around barriers include CGCG, as well as AAAA, here seen with its characteristic asymmetric distribution mirrored by the opposite pattern for TTTT. TATA, which can be a core promoter element, shows a slight underenrichment near the identified barriers.

To explore this, we selected the region around the *RER2* gene, which has two domain boundaries according to Micro-C data (Figure 6(b)). We systematically injected an *Abf1* motif at every base-pair position in this region, predicted the perturbation effect (wild-type - mutant) and quantified the lean effect. When inserted slightly to the left of the first barrier, the perturbed nucleosomes lie to the left of the motif, while insertion slightly to the right produced right-leaning effects (Figure 6(c)).

Interestingly, when these perturbation effects were visualized for every position, they resemble Micro-C interaction maps, either from genome-wide *in vivo* data[71] (Figure 6(d)) or from *in vitro* data on a targeted region[73](Extended Data Figure 8(a)). Thus, we identified previously mapped boundary elements in intergenic regions at high resolution, here occurring near *Rsc3* and *polyA* motifs (Figure 6(d), Extended Data Figure 8(b)).

To systematically map these barrier elements, we performed motif perturbations across the genome and identified the positions where the lean score flips from left to right. The identified barriers were typically close to *Abf1, Reb1, Rsc3* and *polyA* motifs (magenta lines in bottom of Figure 6(d), Figure 6(e)), suggesting that barriers that resist perturbations by nucleosome-positioning motifs consist themselves of combinations of nucleosome-positioning motifs.

To potentially identify novel motifs, we analyzed the kmer distribution around these barriers (Figure 6(f)). The CGCG kmer, which is part of the *Rsc3* motif, shows the strongest and most central enrichment within 30 bp of barrier elements, while *polyA* kmers are strand-specific and slightly less central. The TATA kmer, which is not directly linked to nucleosome positioning, is underenriched near barriers. We note that the kmer patterns are in closer distances (*σ* = 19 bp) than when we take the boundaries from the Micro-C data[71] (*σ* = 31 bp), suggesting that our method of identifying barriers has higher resolution (Extended Data Figure 8(e)).

We conclude that our model trained on MNase-seq data has learned properties relevant for the 3D organization of the genome. It has learned intricate rules by which motifs position nucleosomes and that motif combinations form barriers that resist the effect of motif perturbations.

### 2.7 BPReveal allows the engineering of sequences with altered nucleosome properties

With the advent of deep learning in genomics, there has been a push to use model predictions to streamline the design of synthetic sequences[74, 75]. To enhance the flexibility of BPReveal’s interpretation tools, we implemented a genetic algorithm (GA), to mutate sequences in a way that maximizes a desired experimental outcome, defined and bounded by the user. We used the GA to change the nucleosome configuration on the well-characterized *Pho5* promoter, which is bound and induced by the TF Pho4 under phosphate starvation[76, 77](Figure 7).

**Fig. 7.**
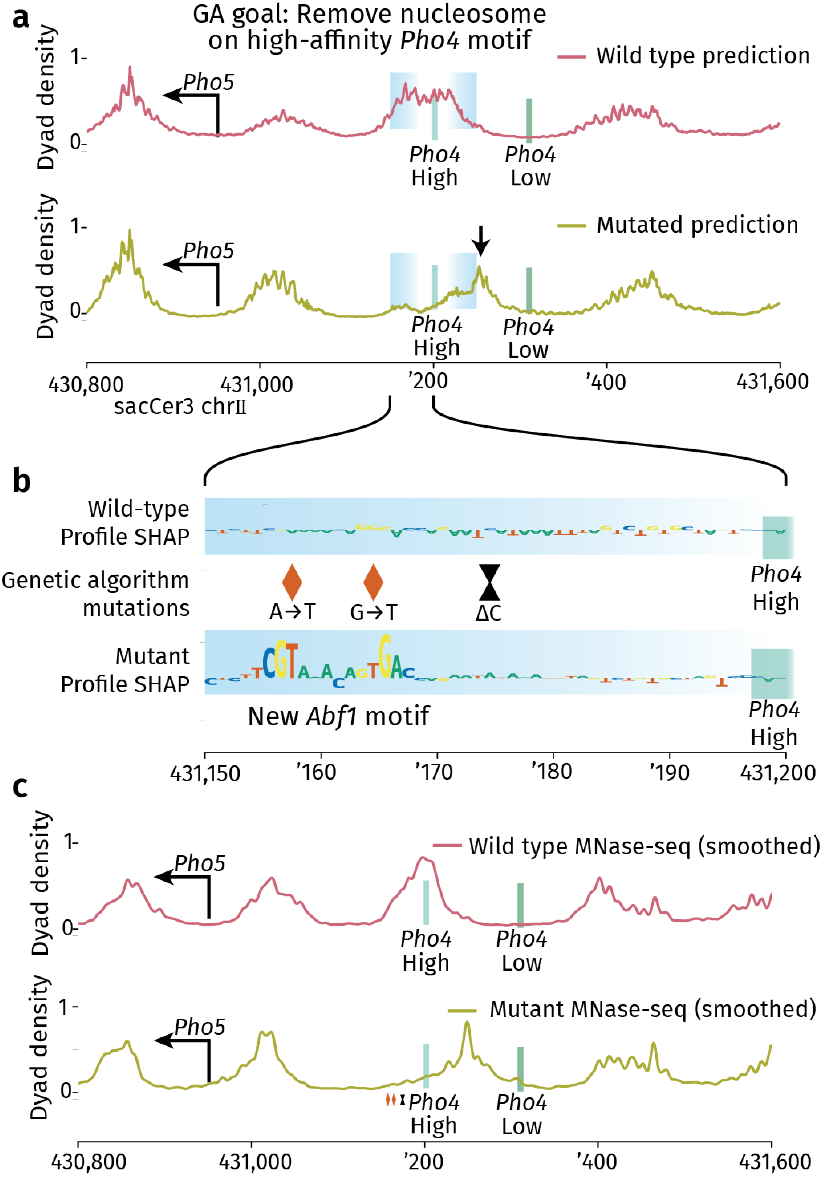
BPReveal’s genetic algorithm (GA) generates sequence designs for nucleosomes that can be experimentally validated. (a) The GA feature in BPReveal was directed to reduce nucleosome occupancy in the blue region centered on the high-affinity *Pho4* motif with three mutations. It returned mutations that induce a predicted shift of the nucleosome to the right side (indicated with an arrow). (b) The mutations introduced a new *Abf1* motif (orange diamonds) about 20 bp away from the *Pho4* motif, as visualized by the contribution scores obtained with DeepSHAP. An additional mutation (black hourglass) has only small effects. (c) Experimental validation of this sequence design was performed by CRISPR/Cas9-mediating editing of the endogenous yeast PHO5 locus and performing MNase-seq on the wild-type and mutant strain (shown smoothed as there is no bias correction). This confirmed that the nucleosome is shifted from the *Pho4* motif, while leaving the rest of the nucleosome landscape largely unperturbed.

Under repressed conditions (i.e., high phosphate), a low-affinity *Pho4* motif is exposed, while a high-affinity *Pho4* motif is covered by a nucleosome (Figure 7(a))[78–80]. Since previous studies have analyzed the consequences of mutating the two *Pho4* motifs[81], we decided to manipulate the nucleosome configuration without perturbing the motifs.

We used the GA to create a sequence design that minimizes the nucleosome on the high-affinity *Pho4* binding site with a maximum of three mutations. We obtained a design where two of the designed mutations introduce a new *Abf1* motif, which is predicted to create a nucleosome-depleted region and expose the *Pho4* motif by shifting the nucleosome to the right (arrow in Figure 7(a)). A third mutation only slightly enhanced the effect (Figure 7(b)). By repeating the GA sequence search, we found that the introduction of TF motifs was a frequent solution. Of the 409 different runs, 77 of them introduced a new *Abf1* motif, and 122 introduced a *Reb1* motif.

To validate the GA’s sequence design, we created a yeast strain where we introduced the three mutations via CRISPR/Cas9-mediated editing. We then performed MNase-seq to map the nucleosomes in the wild-type strain and the mutant strain (Figure 7(c)). We found that the nucleosome that covers the *Pho4* binding site in wild-type is shifted to the right in the mutant, thereby exposing the high-affinity *Pho4* site, as we had intended (Figure 7(c)).

These experimental results demonstrate that our nucleosome model allows the design of strains with altered nucleosome properties, which represents an opportunity for simultaneously studying the role of DNA sequence, nucleosome positioning, and gene regulation in the future. Taken together, this highlights the versatility of BPReveal in predicting and interpreting a variety of genomics data sets, and the use of models to create synthetic sequence designs.

## 3 Discussion

PISA is, at its core, a way to ask how one stretch of DNA affects a biological signal in its surrounding region. We are not the first to ask this question, as classical genetic screens and statistics-based approaches of analyzing genomics data have long provided useful insights[82]. With the recent introduction of interpretable deep learning techniques in genomics, we now have an opportunity to investigate ever-finer positional details of the sequence-to-function relationships that ultimately form the cis-regulatory code. PISA adds to the current set of interpretation tools and has multiple distinct advantages.

PISA works on individual loci in a wild-type context. Traditional statistics-based methods are good at identifying abstract patterns and associations in genomic data, but applying these rules at individual regions of interest has been very challenging. Since deep learning models are trained to make accurate predictions for each region, they inherently learn how to identify and combine multiple sequence elements to predict the experimental outcome in a given region. However, current attribution methods do not fully capture subtle cis-regulatory rules. PISA bridges this gap by providing a precise map by which relevant sequence elements drive genomics readouts at single-base resolution.

PISA plots are inherently visual and intuitive. We have used PISA heatmaps and squid plots here for four different experimental datasets from three different organisms, and continue to find them useful in many contexts. The visualizations highlight the range and patterns by which motifs affect output predictions, and we could often link these patterns to known mechanisms. For example, in a model of TF binding in mESCs, PISA revealed a different influence range of the *Oct4-Sox2* and *Nanog* motif, consistent with their distinct effects on chromatin accessibility[35]. We also discovered a strong, localized footprinting effect on chromatin accessibility by a CA-repeat motif, *CAckle*, which is obscured when using traditional attribution methods but known to be functional in transcription studies[13, 39]. Finally, we observed a leaning motif effect of nucleosome-positioning effects in PISA plots, which inspired our investigation into chromatin domain boundaries.

PISA provides an easy way to identify irrelevant patterns such as experimental biases. Enzymatic biases in genomics assays are typically mixed in with the biological signal, and thus hard to distinguish with traditional attribution methods. In PISA heatmaps, the contribution scores of local enzymatic biases are distinguishable as distinct diagonal bands. In the case of MNase-seq, we leverage this property to create an experimental bias track, which in turn can be used to train a bias-minimized MNase-seq model.

The ability to train a bias-minimized MNase-seq model from sequence alone and discover TF motifs that contribute to the nucleosome organization is unprecedented. Using yeast data, we discovered and visualized the long-range nucleosome positioning effects of motifs, identified these motifs as domain boundaries in the 3D chromatin organization, and designed minimal mutations that precisely reposition nucleosomes *in vivo*. Our approach to identify barriers is complementary to those using Micro-C data, but has higher resolution and considerably lower sequencing depth requirements for larger genomes. We expect this approach to be applicable to more complex organisms, as long as the MNase-seq data are of high quality and high coverage.

Taken together, BPReveal and PISA provide a useful method for training a variety of experimental data and visualizing the intricate rules of the cis-regulatory code across various species. The approach is particularly useful for mapping and understanding the sequence basis of nucleosome positioning, enabling us to better study the role of nucleosomes in the cis-regulatory code.

There are limitations of PISA that should be kept in mind. First, PISA plots are only as good as the underlying model, and thus careful performance benchmarking is required before investing into interpretation. Second, PISA reveals sequence rules that the model learned, but not the proteins and regulatory mechanisms that create these rules inside cells. For example, PISA revealed that the *Zelda* motif not only drives chromatin accessibility but also the acetylation on the flanking nucleosomes. Supported by previous data[15, 83], we can guess that Zelda interacts with an acetyltransferase, but we cannot rule out that the effect is mediated indirectly through the creation of open chromatin. Such gaps in knowledge are an opportunity for additional experiments, with the long-term goal of linking sequence rules and mechanisms in a coherent framework.

## 4 Methods

### 4.1 Data

#### 4.1.1 ChIP-nexus

A BPReveal model was trained on Oct4, Sox2, Klf4, and Nanog ChIP-nexus experiments in mouse R1 mESC lines using the same processed bigwig tracks and peak coordinates as in the original study (GSE137193[84, 85]). To the extent possible, we maintained the same model parameters as used previously.

#### 4.1.2 ATAC-seq

A ChromBPNet-style model[14] was trained on ATAC-seq data from 2-3h *Drosophila melanogaster* embryos using the BPReveal framework. The same processed bigwig files, peak coordinates, and model parameters were used for training (GSE218852[15, 86]).

#### 4.1.3 Published MNase-seq data

We used a published MNase-seq data set, specifically the wild-type experiments SRR12073988 and SRR12073989 from GSE153035[48]. Paired-end reads were aligned against the sacCer3 genome using bowtie2 `--very-sensitive -X 1000` [87] (Bowtie2 version 2.5.1). Aligned fragments that spanned more than 1 kb were eliminated, but no other size selection was performed. For all paired reads, we created coverage tracks of the 3^*′*^ endpoints, 5^*′*^ endpoints, and fragment midpoints. (3^*′*^ and 5^*′*^ in this context are with respect to the genome; therefore all fragments are effectively treated as being on the positive strand.) The code used for this processing is included in the code repository for this paper at https://github.com/zeitlingerlab/bpreveal-manuscript.

For the performance comparison against Routhier *et al*[25], we used the tracks in the GitHub repository for that paper.

#### 4.1.4 Micro-C data

We used the Micro-C data from Hsieh et al.[71], with GEO accession GSE68016. Specifically, we used the wild-type replicates with accession numbers GSM1661107 to GSM1661126. Data were aligned to the sacCer3 genome using `bowtie2 -U` `--very-sensitive-local`, and reads were selected for plotting using custom scripts which are available in the code repository for this manuscript. Briefly, all paired reads entirely contained in the display window were used to build the contact map, and each pixel in the contact map represents 50 bases.

To compare our identified barriers to those identified with Micro-C, we used the set of boundaries published previously[71].

#### 4.1.5 Histone modification ChIP

We used published H3K27ac ChIP-seq data from[15, 86], specifically from the GSM6757761, GSM6757762, GSM6757763 and GSM6757764 datasets. We used the same pipeline to call peaks and process bigwig files.

#### 4.1.6 Custom yeast mutants

The *S. cerevisiae* strain BY4741, a derivative of S288C with the genotype MATa his3Δ1 leu2Δ0 met15Δ0 ura3Δ0 was used. A single isolate was whole-genome sequenced before strain construction. The three point mutations were introduced using CRISPR/Cas9-based genome editing as described previously[88]. A 20 bp gRNA close to the mutation site was designed with BsaI overhangs and ordered as oligonucleotides. The gRNA-encoding oligonucleotides were annealed and cloned into the pCASB plasmid (Addgene 190175) using BsaIHF-v2 enzyme and T4 DNA ligase. The cloning reaction was transformed into *E. coli* and plated on Kanamycin selection plates. The plasmids were then isolated and verified by Sanger sequencing. A 160 bp homology-directed repair (HDR) template was designed to contain the desired mutant sequence, synthesized by Genescript and amplified by PCR. Yeast cells were then co-transformed with the cloned plasmid encoding Cas9 and gRNA, along with the HDR template. Transformants containing pCASB were selected on G418-containing media. Colonies were streaked twice, and isolates were screened for loss of the gRNA plasmid via replica plating to G418. For each isolate, genomic DNA was extracted, and the region of interest was amplified by PCR, purified, and verified by Sanger sequencing.

#### 4.1.7 MNase-seq experiments

MNase-seq experiments were performed essentially as described previously[89] in replicates. Yeast cultures were grown at 30°C in YPD to OD_600_ 0.8-1 and crosslinked with 1% formaldehyde for 15 minutes at room temperature. 125mM Glycine was added to quench the reaction. Cell pellets were resuspended in spheroplasting buffer (1M Sorbitol, 5 mM *β*-mercaptoethanol, 50mM Tris pH 7.5, 2 mg/ml zymolyase (AMS Bio cat no: 120493-1); 1 mL of buffer per 20 mL of cell culture) and incubated for 15 min. at room temperature. The derived spheroplasts were treated with 100U MNase (NEB cat no: M0247S) for 30-40 min at 37°C. The reactions were stopped by the addition of EDTA pH 8.0 (50 mM final conc.) and EGTA pH 8.0 (50 mM final conc.). Samples were then incubated with RNase A (Thermo Scientific, EN0531, final conc. 0.2 mg/ml) at 42°C for 30 min to digest RNA. The crosslinks were reversed by adding SDS and proteinase K (Invitrogen cat no: 25530049, 1mg/ml final concentration) and incubation at 65°C for 45 min. DNA was extracted using the Monarch PCR & DNA cleanup kit. Samples were resolved on 1% agarose gel to evaluate the digestion. Mononucleosome-sized bands were extracted and libraries were constructed from 10ng purified DNA using the Watchmaker DNA Library Prep kit (cat no. 7K0102-096) from Watchmaker Genomics according to the manufacturer’s instructions. Paired-end sequencing was performed on AVITI (2x 75bp cycles). The data processing was identical to that used for the published MNase-seq samples.

### 4.2 BPReveal implementation

The BPReveal package is a suite of tools for flexible training and interpretation of BPNet-derived sequence-to-profile models[3, 9, 14]. The code base is licensed under the GNU General Public License (Version 2 or later) and is available at https://github.com/mmtrebuchet/bpreveal. The architecture is shown in Extended Data Figure 1. The input is a one-hot encoded DNA sequence, with typical input lengths around 3,000 bp. The models consist of an initial convolutional layer, followed by a stack of dilated convolutions that exponentially increase in receptive field. There are two outputs for each dataset: (1) the “profile” output is a vector representing the log-probability (i.e., logits) of finding a read at each position in the output window; (2) the “counts” output is a scalar representing the natural logarithm of the total number of reads observed in that window. For brevity, the two outputs are referred to as one “head” per dataset. The model can be trained as a multi-task model on several experiment types simultaneously, including but not limited to ChIP-seq, ATAC-seq, MNase-seq, and PRO-cap data.

BPReveal includes two significant expansions from the original BPNet architecture that were first implemented in ChromBPNet [14] and ProCapNet [9]. First, instead of giving convolutional filters beyond the input window zeros, as is often done in image processing, the input length is increased to provide DNA sequence for the entire receptive field [14]. Second, multiple strands from one experiment can be combined into one output. The log-counts output then combines the total reads from both strands, and the profile logits have the shape (`output-length, num-strands`). This benefits the training of data such as PRO-cap, where reads are often primarily on one strand [9].

The loss is the same as it was in the original BPNet:

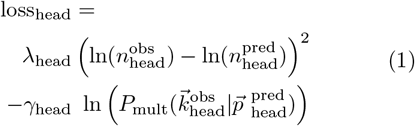

and

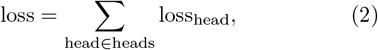

where *γ*_head_ and *λ*_head_ are (user-assigned) weights to the profile and counts components of the loss for each head, ln 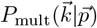 is the log-likelihood of observing the outcome 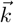 (i.e., the observed experimental reads) given the probability distribution 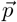 (i.e., the predicted logits from the model), and *n* is the total number of reads observed (or predicted) over an entire region.

### 4.3 Bias regression

BPReveal includes a tool based on ChromBPNet[14] to remove experimental biases from genomics data. The technique used in ChromBPNet was based on the assumption that

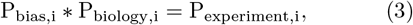

where P_bias,i_ represents the probability density at base *i* due only to experimental biases, P_biology,i_ represents the probability density of the true biological profile at base *i*, and P_experiment,i_ is the probability density that would be measured by an experiment. Since BPReveal and ChromBPNet models both use logits to represent the output profiles, this becomes

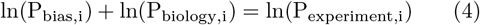

in the implementation.

BPReveal implements this strategy using two BPNet-like models: One submodel, called the solo model, is pre-trained to predict just the experimental bias and its weights are frozen during the training of the second submodel. The other submodel, called the residual model, has its output logits added to the output logits of the frozen bias model. These combined logits are then used in the standard loss function, Equation 2.

A feature introduced by BPReveal is that the bias is transformed to better match the experimental data before its weights are frozen:

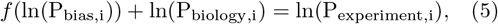

where *f* is a simple function applied to all of the outputs from the bias model. For all of the models used in this paper, we used the function *f*(*x*) = *w*_1_ * *sigmoid*(*w*_2_*x* + *w*_3_) + *w*_4_. ChromBPNet, by comparison, does not transform the logits from the bias model, and applies a pre-computed scaling factor to the counts output. The full architecture is shown in Extended Data Figure 1. BPreveal also includes features for motif scanning based on the original BPNet algorithm[3].

### 4.4 PISA

The calculation of PISA values ℙ_*i* → *j*_ for a particular genomic coordinate *j* is implemented in BPReveal in the following way. An input sequence is selected such that the prediction for base *j* will occur in the output of the model’s predicted profile. deepSHAP is then used to partition the logit at position *j* among each input base *i* that is in the receptive field of base *j*. The references used for deepSHAP are generated by shuffling the input sequence; an option is provided to perform kmer-preserving shuffles using the `ushuffle` algorithm[90].

In the following pseudocode, `i` and `j` refer to *genomic* coordinates, and so `shapVals[0]` refers to ℙ_(*j*− buf) →*j*_. For ease of calculation, we choose input sequences such that base *j* is at the leftmost output of the model.

~~~
buf = (inputLength - outputLength) // 2
sequence = genome.fetch(
  j - buf,
  j + outputLength + buf)
target = model.outputs[headID][0, strandID]
explainer = shap.DeepExplainer(
  (model.input, target),
  ushuffle.shuffleOneHot)
shapVals = explainer.shap_values(sequence)
for i in range(j - buf, j + buf):
 P[i + j, j] = sum(shapVals[i - j + buf])
~~~

This value of the resulting array P holds, at position `(i,j)`, the value of ℙ_*i*→*j*_.

### 4.5 Synthetic bias calculation

The ChromBPNet approach to removing experimental bias has a key limitation: it requires a model that has been trained to predict only bias. This is not straight-forward for MNase-seq data. Therefore, a synthetic bias track for MNase-seq is derived using PISA in the following way.

Consider an ideal dataset giving the exact positions of nucleosomes by their 5^*′*^ and 3^*′*^ end points, with no experimental bias. These data could be used to train a two-strand model *I*; this model would have outputs 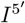 and 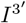. Consider a nucleosome that spans from base 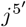 to 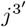. If a base *i* contributes to this nucleosome’s position, then

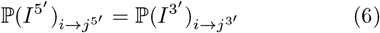

where 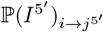 represents the PISA contribution identified by 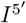 from the base at position *i* (the x-axis in a PISA heatmap) onto the readout at position 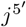 (the y-axis in the PISA heatmap). Because base *i*’s role to causing the 5^*′*^ endpoint to occur at base 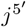 is the same as its role in causing the 3^*′*^ endpoint to occur at base 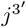, these two contributions will be equal. (This assumption is not strictly true in the case of a partial nucleosome, where a base might not alter the position of a nucleosome dyad but would cause only one observed endpoint to shift. We have found that these instances are rare enough in the genome to not affect our synthetic bias model.)

Now consider the case of real MNase-seq data that has been used to train model *M*. In this case, the experimental readout is not just based on the nucleosome positions (which would have been captured by the ideal model *I*), but also on the enzyme’s sequence bias that determines the observed endpoints. We refer to a model that captures this enzymatic bias as *B*. By Equation 5, we have *f*(*P*_*B,j*_) * *P*_*I,j*_ = *P*_*M,j*_ for each base *j*. 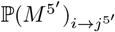 still includes base *i*’s role in causing a nucleosome to have its 5^*′*^ endpoint at base 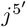, but it also captures the role of base *i* in whether or not the MNase enzyme would stop at base 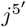 due to bias. This introduces an asymmetry: by the nature of the MNase enzymatic bias, however, base *i* will only have an effect on the enzyme’s bias at 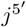 if *i* is close to 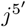. However, 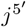 must be about 150 bp away from 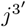 (since they represent opposite ends of the same nucleosome), and therefore if 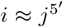 then 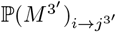 will be due to base *i*’s effect on nucleosome positioning and will have a minimal contribution from bias.

Therefore, for a base *i* close to 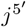:

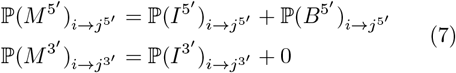

The use of addition here is justified by two points: First, as established by ChromBPNet[14], attribution scores show that addition of logits leads to residual models (i.e., the models without bias) with no contribution from bias motifs. We show in section 2.4 that PISA plots of our MNase residual model show insignificant bias-like density along the diagonal and that our synthetic bias model has density only along the diagonal. Second, since PISA values are (approximate) Shapley values, an effect that is due to a linear combination of two processes will yield Shapley values that are a linear combination of the two corresponding causes: for two models *I* and *B*,

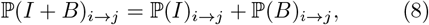

where *I* and *B* are the ideal and bias models, respectively. (and *I* + *f*(*B*) = *M* by Equation 5)

By equations 6 and 7, if 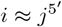 then

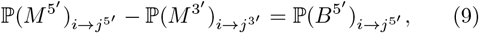

and if 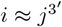 then

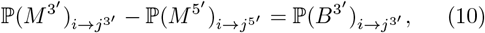

which allows us to calculate ℙ(*B*) from model *M*, which was trained only on the experimental MNase-seq.

In order to calculate the bias at position 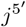 we must know where 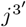 is. We would expect that the offset Δ between the two end points would be on the order of 150 bp (the length of DNA in a nucleosome). We determine the precise value of Δ by comparing the match between 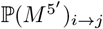 and 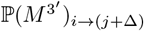 for a range of Δ values. Since the contribution of the bias model should be limited to regions where *i* ≈ *j*, we exclude a 60 bp window around the predicted outputs. In other words, we align the PISA heatmaps of the two strands of model *M* so that they match, except for the places where enzymatic bias is strong. We determined an optimal offset of 179 bp using this method, very close to the average fragment length in our MNase dataset.

By the efficiency property of Shapley values, we can use the bias PISA values to reconstruct a bias profile track:

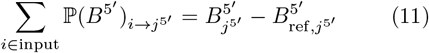

where 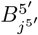 is the enzymatic bias on the 5^*′*^ end of reads starting at base 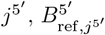 is the output of the model on shuffled copies of that same sequence, and input contains all bases that are in the receptive field of the model when it makes a prediction at base 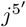. The average enzymatic bias on many random sequences 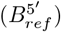 will be uniform, and since 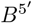 predicts logit values, a constant offset has no effect on the profile. (Equation 11 is formulated in terms of the 5^*′*^ bias, but the same algorithm also applies to the 3^*′*^ case.) Therefore, by summing the rows of the synthetic bias PISA heatmap, we can derive a profile of MNase bias.

The essential process of shifting and subtracting is given in the following pseudocode:

~~~
def getBias(P3, P5, strand):
  # P3 and P5 are the PISA heatmaps for the 3’ and 5’
  # endpoints. Strand specifies which head the bias
  # is to be calculated for.
  # Move the 5’ PISA heatmap to the left to align
  # with the 3’ heatmap, or vice versa.
if strand == “3’“:
  P5Shift = shiftLeft(P5, 179)
  difference = P3 - P5Shift
if strand == “5’“:
  P3Shift = shiftRight(P3, 179)
  difference = P5 - P3Shift
 # The bias will come from near the diagonal, so zero
 # PISA values more than 13 bp from the diagonal as
 # we know they are not from bias.
 maxJ, maxI = difference.shape
 for i in range(maxI):
  for j in range(maxJ):
   if abs(i - j) > 13:
     difference[j,i] = 0
# Use the efficiency property of Shapley values
# to generate a bias track.
biasLogits = sumRows(difference)
bias = softmax(biasLogits)
# Bias has shape (maxJ,)
return bias
~~~

A full implementation can be found in `extractBiasBigwigs.py` in the repository for this paper.

To get a genome-wide profile of MNase bias, we performed genome-wide PISA. In *S. cerevisiae*, this takes a few days on three Nvidia A100 GPUs. For a significantly larger genome, performing PISA on the entire genome is impractical. But since we train a model on the synthetic bias track, it is only necessary to produce enough synthetic bias data to train a model, and then that model can predict the bias genome-wide.

With this synthetic bias track in hand, a model can be trained on pure bias, which is then used to train a residual model à la ChromBPNet, thus learning the underlying biology that mixes with the bias to give rise to the experimental output.

### 4.6 Lean values and boundary identification

To identify barriers from a trained MNase-seq model, we systematically insert a nucleosome-positioning motif at every position *k* across the genome (or across regions of interest). For each motif insertion, we then calculate the lean value *L*^*k*^, which records whether the insertion at position *k* tends to perturb nucleosome positions more to the left or to the right. Barriers are the positions *k* where the lean values *L*^*k*^ transition from positive to negative.

For robustness, we performed the procedure with four different nucleosome-positioning motifs: *Abf1* (ATCATTGTGCACGG), *Rsc3* (`CGCG`), *Reb1* (GCCGGGTAAC), and *polyA* (`AAAAAAA`).

To obtain lean values *L* for each position *k*, the motif is inserted in the center of the output window of length *N* (the prediction window is shifted for each motif position *k*). We then calculate the mutation effect 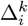 at each position *i* of the output window:

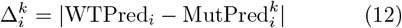

where WTPred and Mutℙred^*k*^ are the predicted MNase profiles (midpoints, not 3^*′*^ or 5^*′*^ end points) for the wild-type sequence and the mutated sequence with the motif insertion at position *k*. WTℙred, Mutℙred^*k*^ and Δ^*k*^ are vectors of length *N*.

Then we calculate the sum of the differences on the left side 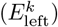 and right side 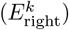 of the injected motif at position *k*. Both are scalars with reads as units:

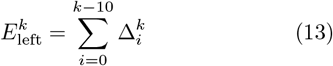

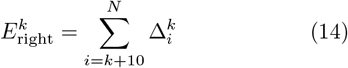

The lean value *L*^*k*^ (also a scalar) is then defined by:

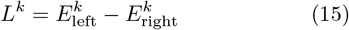

*L*^*k*^ is positive when the effect is primarily to the left of the injected motif and negative when the effect is to the right. After performing this calculation genomewide, we have a value of *L* at each position in the genome (except for chromosome edges).

To identify barriers in the lean values, we applied two orthogonal methods to detect transitions from positive to negative lean values. The first method is based on counting positive and negative lean values. For a position *k*, if the lean values to the left of *k* are mostly left leaning and the lean values to the right of *k* are mostly right leaning, then position *k* represents a point where a motif’s influence flips from one domain to another:

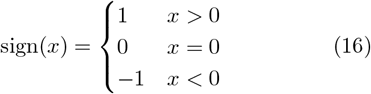

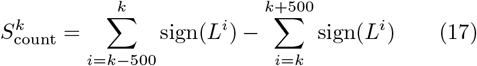

When *S*_count_ is maximal, the lean values flip from mostly positive to mostly negative, thus representing a candidate barrier.

Our second metric for identifying barriers, *S*_deriv_, is based on the derivative of the lean values. When the derivative is large and negative, the lean values are rapidly switching from left to right, potentially representing a barrier. To avoid tracking the nucleosome-scale oscillation observed in lean values, we added *B* as a filter:

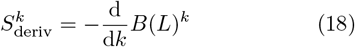

where *B* represents a 10th-order bidirectional Butterworth lowpass filter with a critical frequency of 1*/*200. When *S*_deriv_ is maximal, the lean values are rapidly decreasing, pointing to a candidate barrier.

To identify the maxima in the two score tracks *S*_count_ and *S*_deriv_, we applied `scipy.signal.find_peaks`. The `height` parameter was set to either the 90th percentile (derivative) or 70th percentile (counting) of the corresponding scores, the `distance` between peaks was at least 200 bp, and the `prominence` was set to twice the `height`.

To derive reproducible boundary elements from these candidate barriers, we combined those from both scores, *S*_count_ and *S*_deriv_, for each of the four inserted nucleosome positioning motifs motifs. Boundary elements were called if they scored in at least five out of the eight tests. The lean value tracks, along with the called barrier elements, are available in the `insulators` directory of the data deposited with this paper.

To calculate the enrichment of kmers around barriers, we identified all matches to the kmer or motif within 200 bp of all barriers, and then mean normalized the resulting histogram. To determine the spread of the distribution, we used `scipy.optimize.curve_fit` to match a Gaussian function with a constant offset:

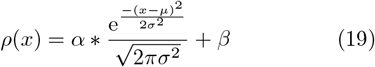

where *ρ*(*x*) represents the frequency of occurrence of a particular motif or kmer at a distance *x* from a barrier. *µ* is the location of the peak relative to the barrier, *σ* is the standard deviation of the Gaussian function, *α* is an arbitrary scaling constant, and *β* is a constant offset to match the genomic background content of the motif or kmer.

### 4.7 Genetic algorithm

To design novel sequences with a desired profile, we implemented a genetic algorithm (GA). This GA designs small sets of mutations that can be applied to an initial sequence in order to maximize a userdefined property of the prediction. Instead of operating directly on the sequence, our GA is given a pool of possible mutations (usually, every substitution, insertion, or deletion, with exceptions to prevent important sequence features from being altered). The optimization procedure then selects a subset (of user-defined size) of the possible mutations that maximize the fitness function. In this way, the algorithm can explore a wide array of possible mutations, including indels, but the edit distance from the WT sequence is limited to the size of the set of mutations. To design the mutation presented in Figure 6, we allowed for three mutations and our fitness function minimized the nucleosome density in a window spanning chrII:431150-431250. Since we expected to test our mutations *in vivo* by using CRISPR/Cas9-mediated editing, we ran the GA once for every possible PAM site in the region, allowing mutations up to 50 bp downstream of the PAM site. We disallowed mutations inside the *Pho5* gene body, on the *Fkh2* motif, or on either of the *Pho4* motifs. Of the 51 runs (one for each PAM site), we manually selected a design that was predicted to remove the nucleosome on the high-affinity *Pho4* motif while leaving the other nucleosomes undisturbed.

### 4.8 Sequence-based motif scanning

To identify motifs purely from sequence, we used Fimo[91] from MEME suite version 5.5.8[82], using the JASPAR databases appropriate for each organism (2022 vertebrates for mouse, 2020 insects for fly, and 2024 fungi for yeast)[92]. For regex-based scanning, we simply searched for instances of the motif in the genome sequence. All relevant code is included in the repository for this paper.

## 5 Author contributions

CEM conceived PISA, developed the software, and performed the analysis. MW, AK, and JZ provided ideas and feedback. JMG constructed the yeast CRISPR strains. GM performed the MNase-seq experiments and analyzed the results. FK optimized parameters for the histone acetylation model. JS provided the PyTorch implementation of PISA. CEM and JZ wrote the body of the manuscript with contributions from all authors. GM wrote the methods for the yeast mutants, and FK wrote the methods for the histone acetylation data processing.

## 6 Acknowledgments

Kyle Weaver and Ben Troutwine provided assistance in generating the CRISPR cell lines used in this work. Alex Garruss, Minal Khatri, and Haining Jiang provided feedback on the manuscript. Jonathon Russell and Joshua Niemeyer provided computational resources and support.

## 8 Data availability

During review, all relevant data, including models, predicted tracks, training data, configuration files, importance score tracks, motifs, and PISA values, are available at https://stowersinstitute-my.sharepoint.com/:u:/g/personal/cm2363_stowers_org/IQAVcwBtXg7URa4WSglPc8KXAer5qkfI5lU1EbTv7dnzfvY?e=ghXOq3. After review, these data will be moved to the Stowers Original Data Repository at https://www.stowers.org/research/publications/libpb-2546. All predicted tracks, models, configuration files, importance score tracks, and motifs are available on Zenodo at zenodo.org/records/15232217 Sequencing data for the *Pho5* site mutations are available on GEO, accession GSE313524.

## 9 Code availability

The BPReveal package is available at https://github.com/mmtrebuchet/bpreveal All source code for this paper is available at https://github.com/zeitlingerlab/bpreveal-manuscript.

## Extended data

**Extended Data Figure 1.**
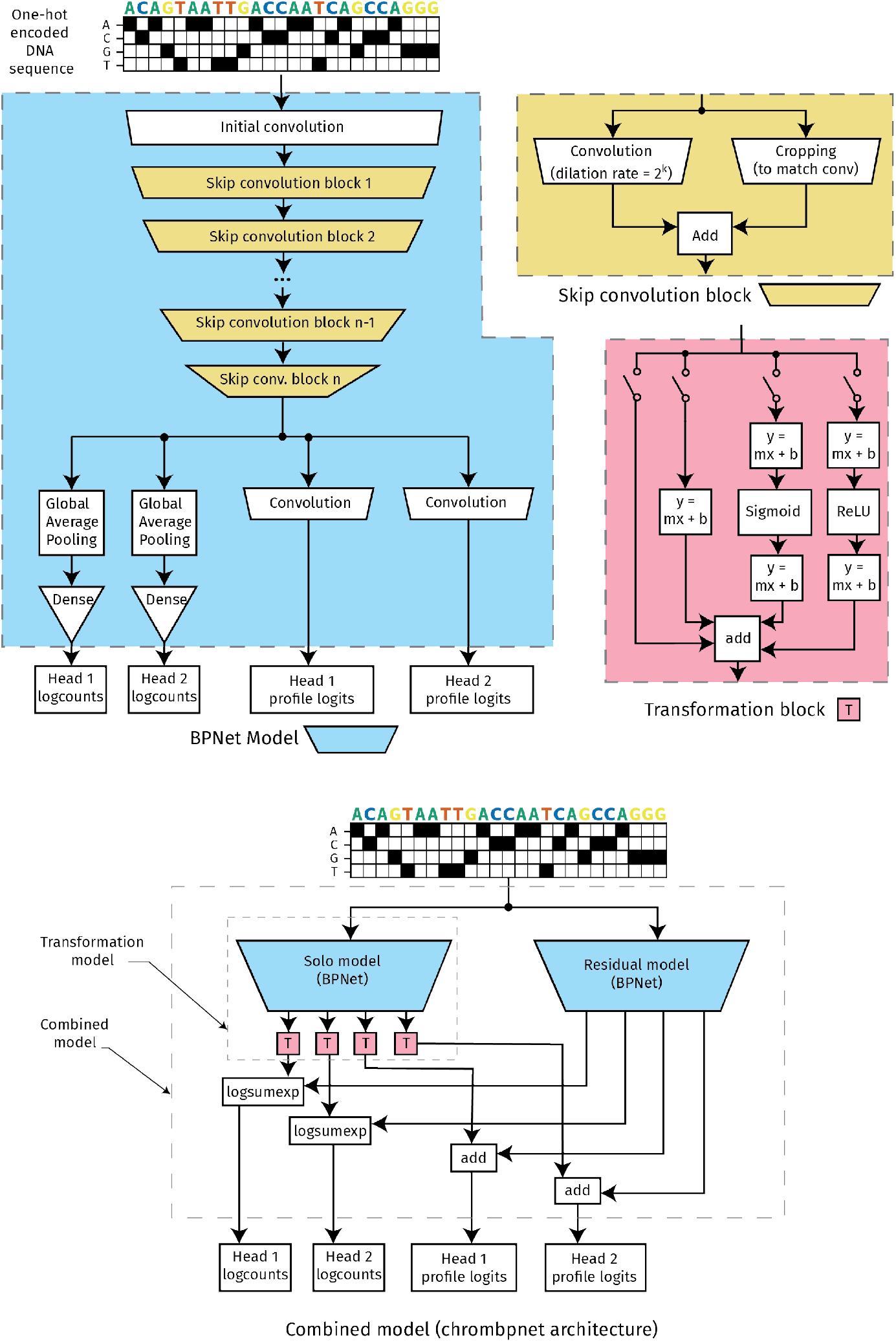
BPReveal architecture. (blue) A typical BPNet-style model with two output heads. The input to a BPReveal model is one-hot encoded DNA sequence. The core of the model is a stack of dilated convolutional layers with skip connections (yellow trapezoids). (yellow) The architecture of a dilated convolutional layer with skip connection. (red) The architecture of the transformation model, used to regress the predictions of a solo (i.e., bias) model to better match experimental data. (bottom) A complete BPReveal model incorporating ChromBPNet-style bias correction. The solo model is pre-trained on enzymatic bias, and then its weights are frozen. One transformation model is applied to each output of the solo model, and the weights in the transformation model are trained on the experimental data. Finally, the weights of both the solo and transformation model are frozen, and a new residual model is trained to learn the experimental data.

**Extended Data Figure 2.**
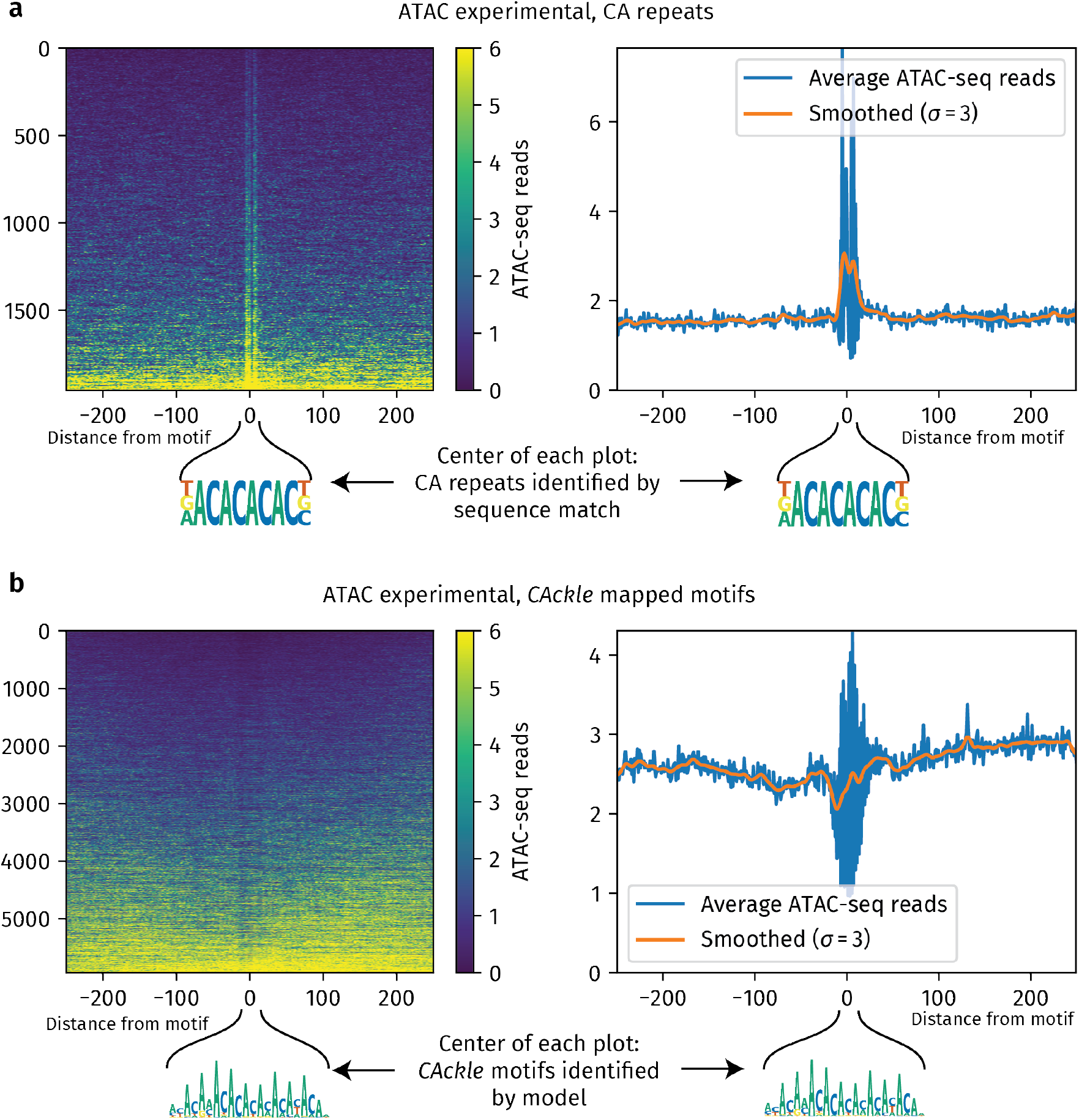
The *CAckle* motif contributes to Tn5 enzymatic bias, but can also create a depleted footprint. We show heatmaps and metapeaks of experimental ATAC-seq data at CA repeats. (a) Mapping CA repeats by sequence alone shows a pronounced spike in ATAC-seq reads at the site of the CA repeat. (left) A heatmap of ATAC-seq profiles at CA repeats, ordered by total ATAC-seq reads in the region. (right) An average profile of the of the CA repeat instances shows that many ATAC-seq reads occur at the site of the repeat. (b) By using model-derived motifs instead of just sequence match, we see that *CAckle* motif instances can cause a local dip in ATAC-seq reads at some regions. A heatmap of ATAC-seq profiles at mapped *CAckle* motifs (left) and an average profile over all mapped motif instances (right) show a slight dip in ATAC-seq reads at the *CAckle* motif. For ease of visualization, a *σ* = 1 Gaussian filter has been applied to both heatmaps. The orange traces in the average profiles are smoothed with a *σ* = 3 Gaussian filter applied to the (unsmoothed) blue average profile traces.

**Extended Data Figure 3.**
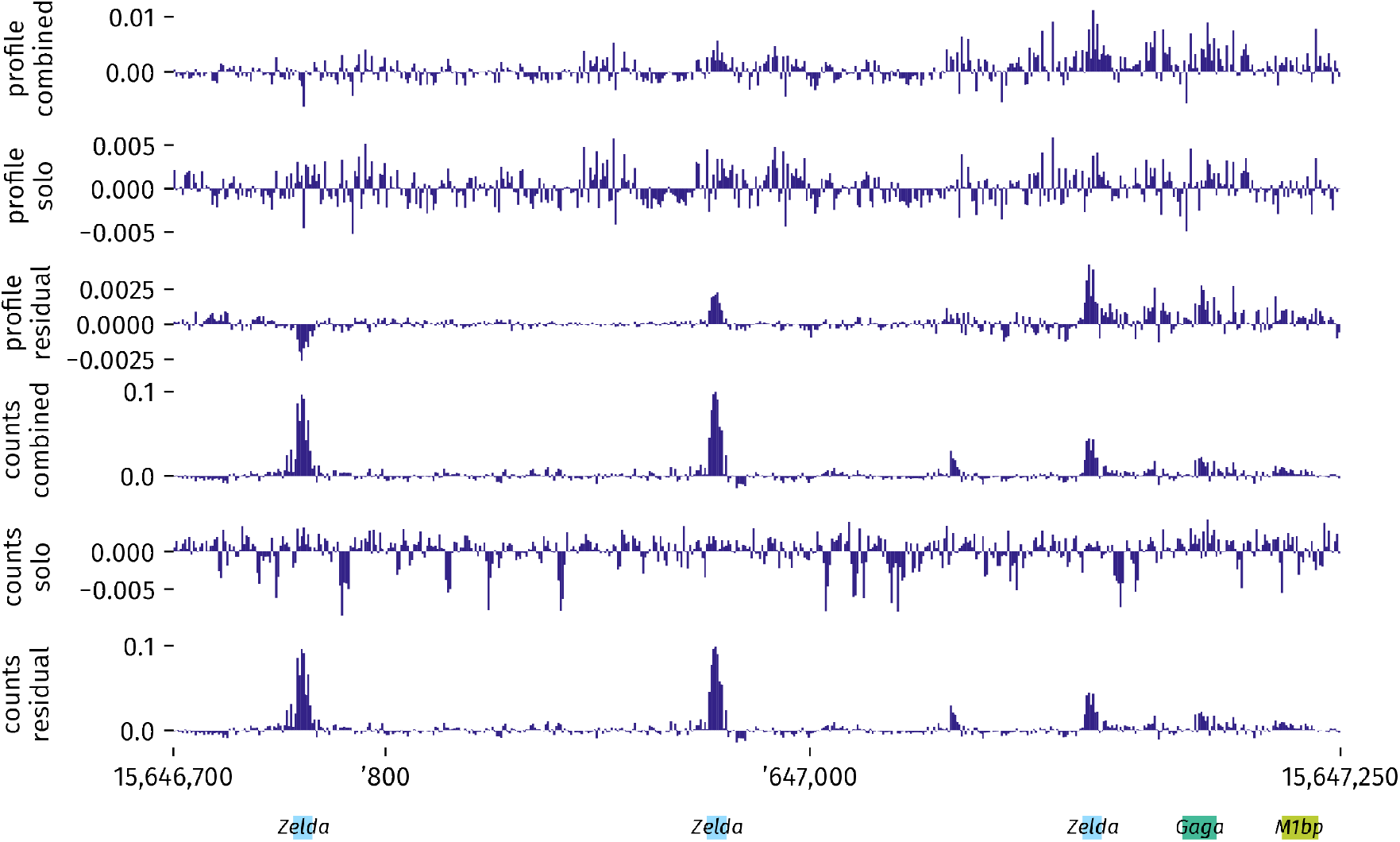
Importance score comparison for ATAC bias correction. The ChromBPNet bias correction strategy leads to a dramatic improvement in profile contribution scores, but does not improve counts contribution scores. We show importance score tracks at the *sog* locus for the combined model (which predicts the actual experimental data, including bias), the solo model (which only predicts bias), and the residual model (which does not include bias). The profile contribution scores for the combined and solo model both show a great deal of noise due to bias, whereas the bias-corrected residual model shows much clearer peaks at the three *Zelda* motifs. The enzymatic bias encountered in ATAC-seq has a much smaller effect on the total counts prediction, such that the combined model already clearly shows the three *Zelda* motifs; the counts contribution scores from the residual model are the same as for the combined model in this case.

**Extended Data Figure 4.**
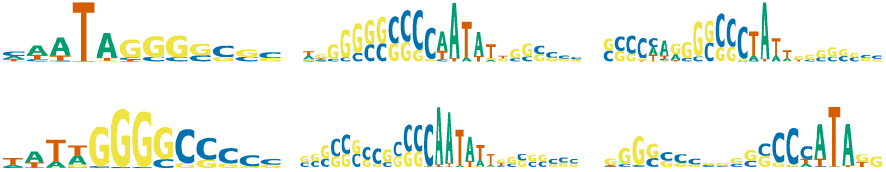
Motifs identified by MNase bias model. Motifs identified from profile contribution scores for our MNase bias model show a transition from AT-rich to CG-rich regions, consistent with the well-characterized bias of the MNase enzyme. The six motifs shown here are the six most frequent seqlets identified by TF-MoDISco; none of the 40 motifs it identified this model resembled a biologically-relevant motif.

**Extended Data Figure 5.**
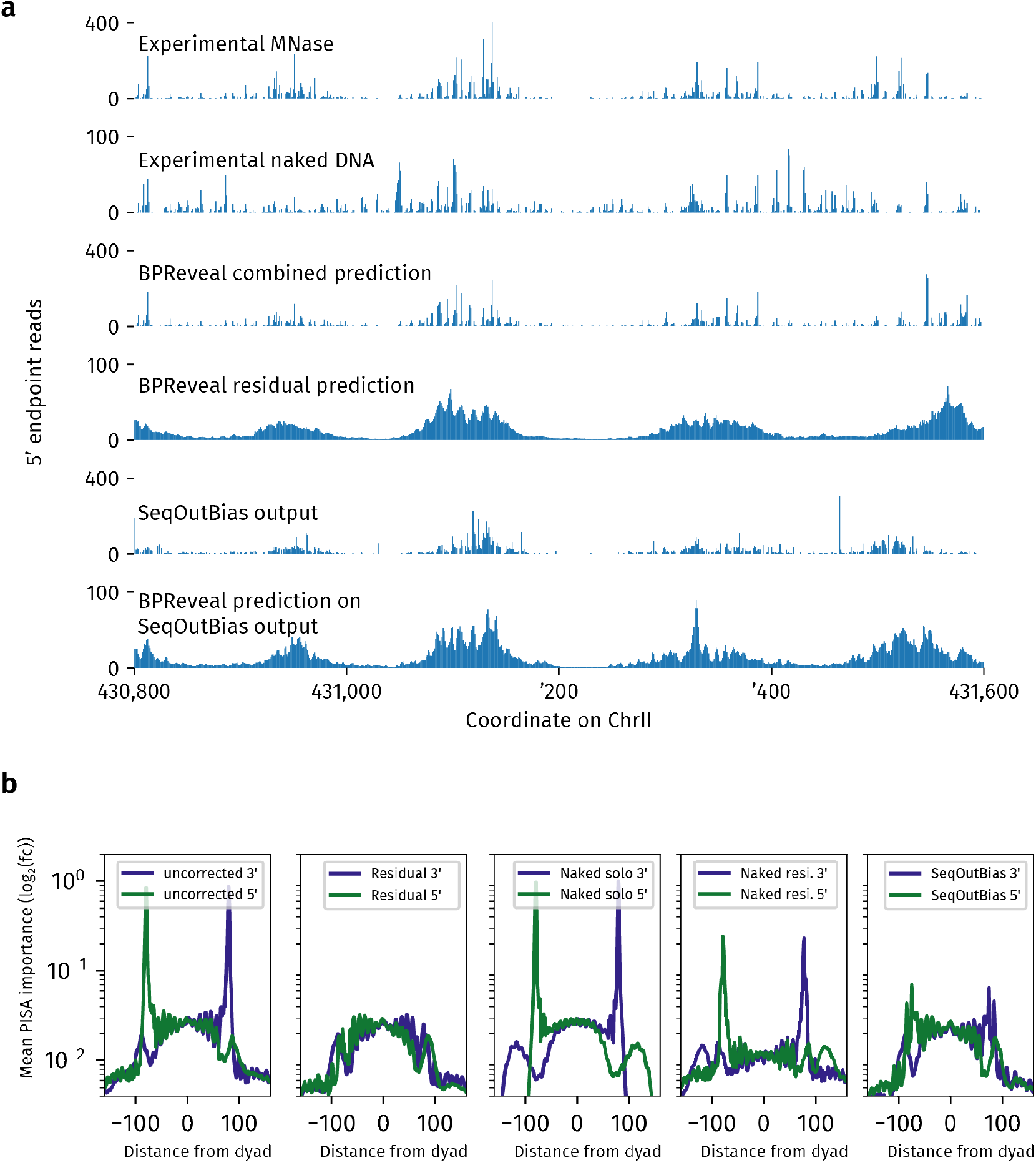
Comparison of other bias-reduction techniques. In (a), we show several tracks that explore potential bias-correction strategies. In each track, we show only 5^*′*^ endpoints of the dataset. Experimental MNase is the MNase dataset from[48]. The experimental naked DNA sample is also from[48] and captures the behavior of the MNase enzyme on naked DNA. The BPReveal combined prediction is a BPReveal model trained on the experimental MNase data, including enzymatic bias. The BPReveal residual prediction is the bias-minimized result using the technique described in this paper. The SeqOutBias output is the track generated by SeqOutBias when given the experimental MNase data. We also trained a BPReveal model on the bias-minimized output from SeqOutBias, which is shown in the bottom track. In (b), we use BAT plots to quantify the effectiveness of bias removal using three techniques. The uncorrected model is trained on MNase data without any bias correction, the residual model uses the bias removal strategy described in this paper, the naked solo model is trained only on the naked DNA experimental track, and the naked residual model is trained using the naked DNA model as a bias model and performing a ChromBPNet-style correction to remove that bias. Finally, the SeqOutBias model was trained on the bias-minimized output from SeqOutBias run on the experimental MNase data.

**Extended Data Figure 6.**
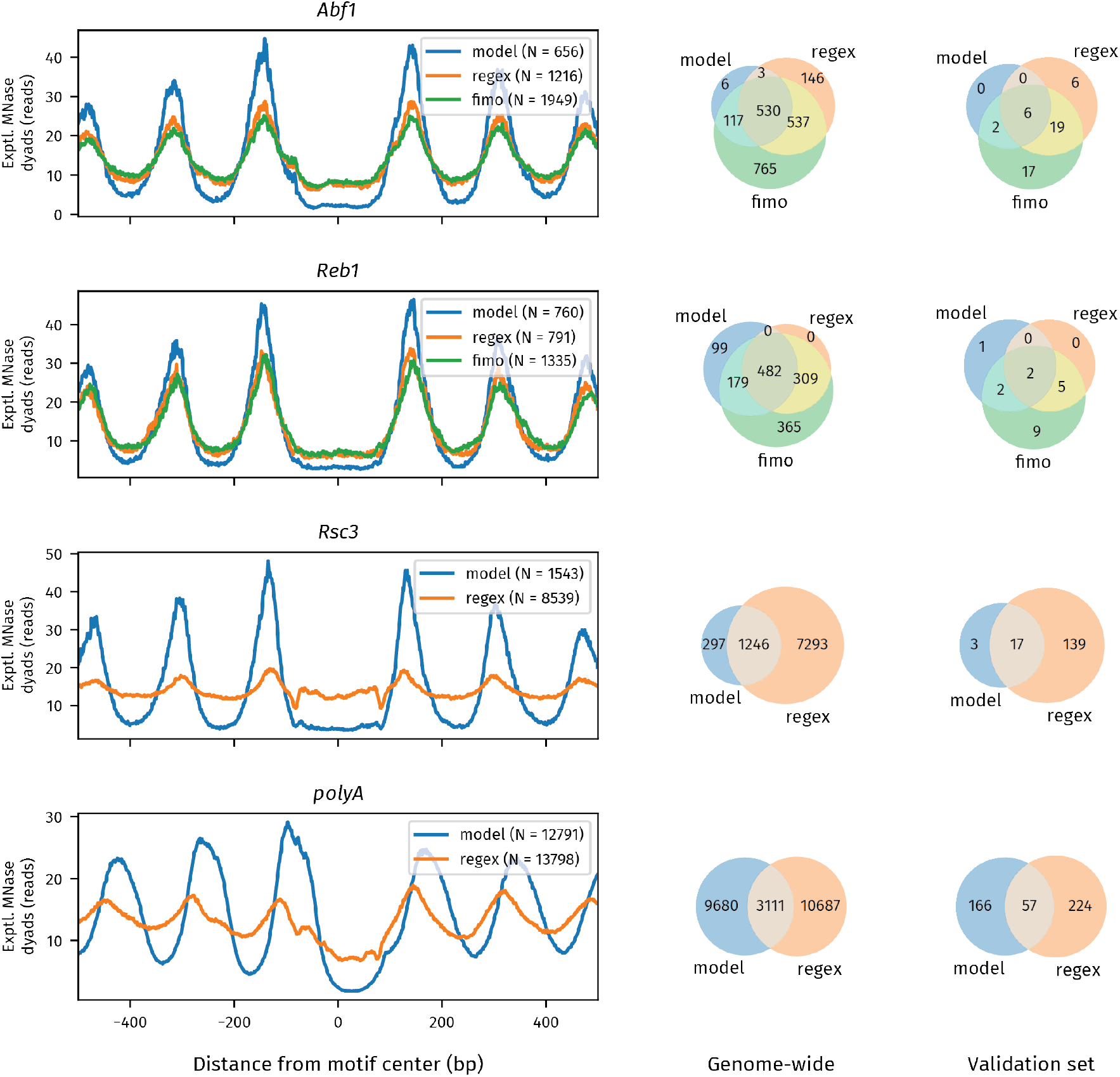
Four motifs identified by MNase residual model. For each motif, we show the average dyad density around that motif when called by three different methods. “model” motifs were discovered by scanning importance scores using the TF-MoDISco patterns. “fimo” motifs are identified using the motif as defined in JASPAR, and are identified purely by sequence match. “regex” motifs are based on scanning the genome to exact matches for a regular expression that matches the motif. The Venn diagram on the right shows how often the motif instances identified by the various methods overlap. For *Abf1* and *Reb1*, the model tends to identify motifs with stronger effects (blue lines) than the text-based algorithms (green and orange lines). For *Rsc3*, even though an *Rsc3* motif is in the JASPAR database, fimo did not identify any instances. The advantage of using a model to identify motifs is evident in that model-derived *Rsc3* motifs have much stronger nucleosome positioning effect than pure sequence-based motif calls. For *polyA*, there is no canonical JASPAR motif, and so fimo cannot be straightforwardly applied. Again, the model is much better at identifying functional *polyA* motifs than a simple sequence match.

**Extended Data Figure 7.**
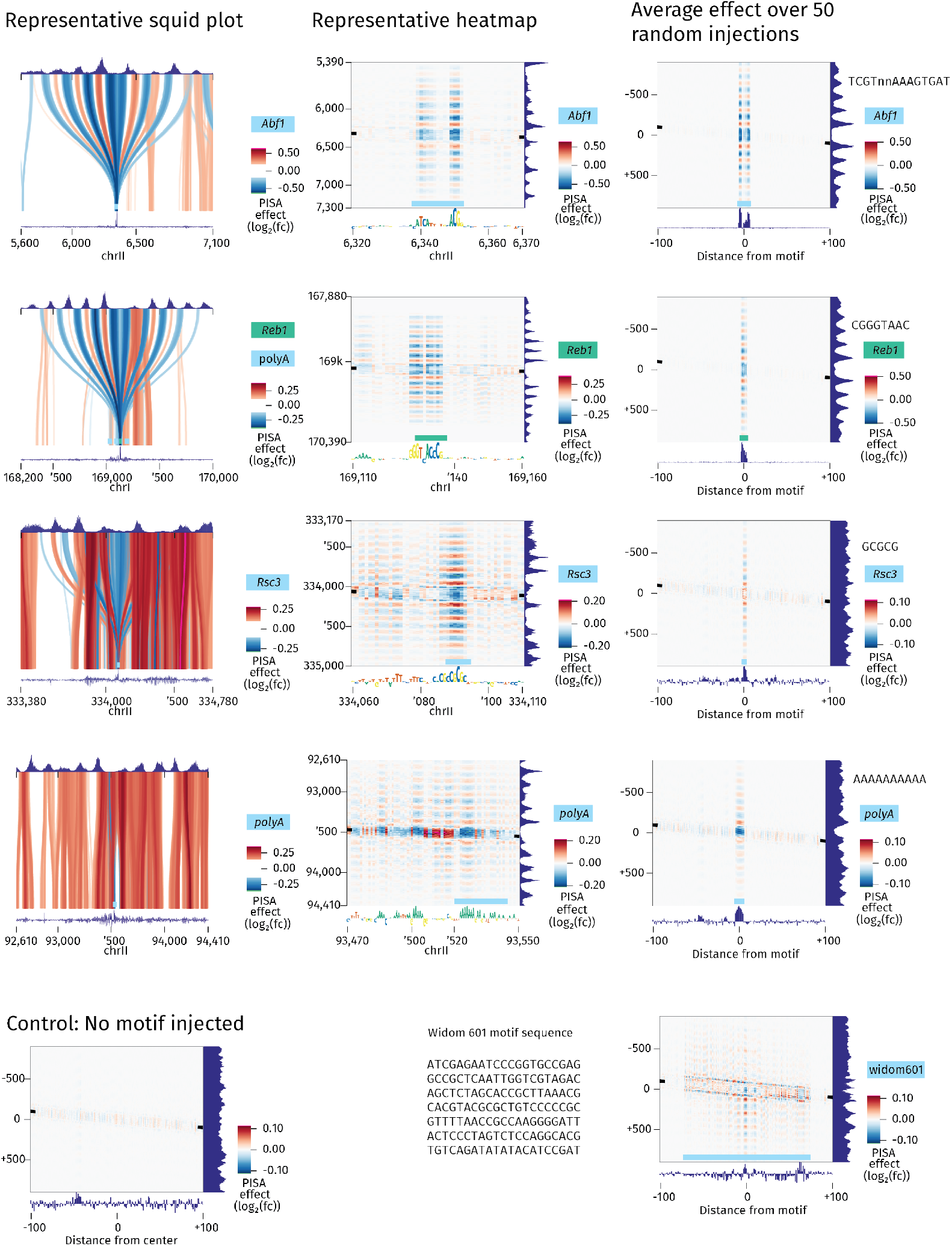
All MNase PISA plots. For completeness, we show PISA squid plots (left) and heatmaps (center) for representative instances of the four motifs discussed in Figure 5 along with average PISA heatmaps generated by injecting the motif in 100 random genomic regions (right). The *Reb1* and *Abf1* motifs both show a characteristic strong positioning effect both in their actual genomic context and when injected into randomly-selected sequences from elsewhere in the genome. The *Rsc3* motif has a weaker effect than the two TFs, but still leads to nucleosome positioning. The *polyA* motif has a more distributed role. While one stretch of A has been identified by our motif mapping tool, several other short stretches of A repeats are also playing a role in nucleosome positioning at this locus. (bottom, left) We show the average PISA heatmaps drawn from 100 randomly-selected genomic regions. All of the injection effects in the right column are considerably stronger than the background level. (bottom, right) The effect of injecting the entire Widom 601 sequence is comparable to the effect of a single *polyA* or *Rsc3* motif injection, despite the Widom sequence being over an order of magnitude longer and specifically designed to position nucleosomes *in vitro*[64].

**Extended Data Figure 8.**
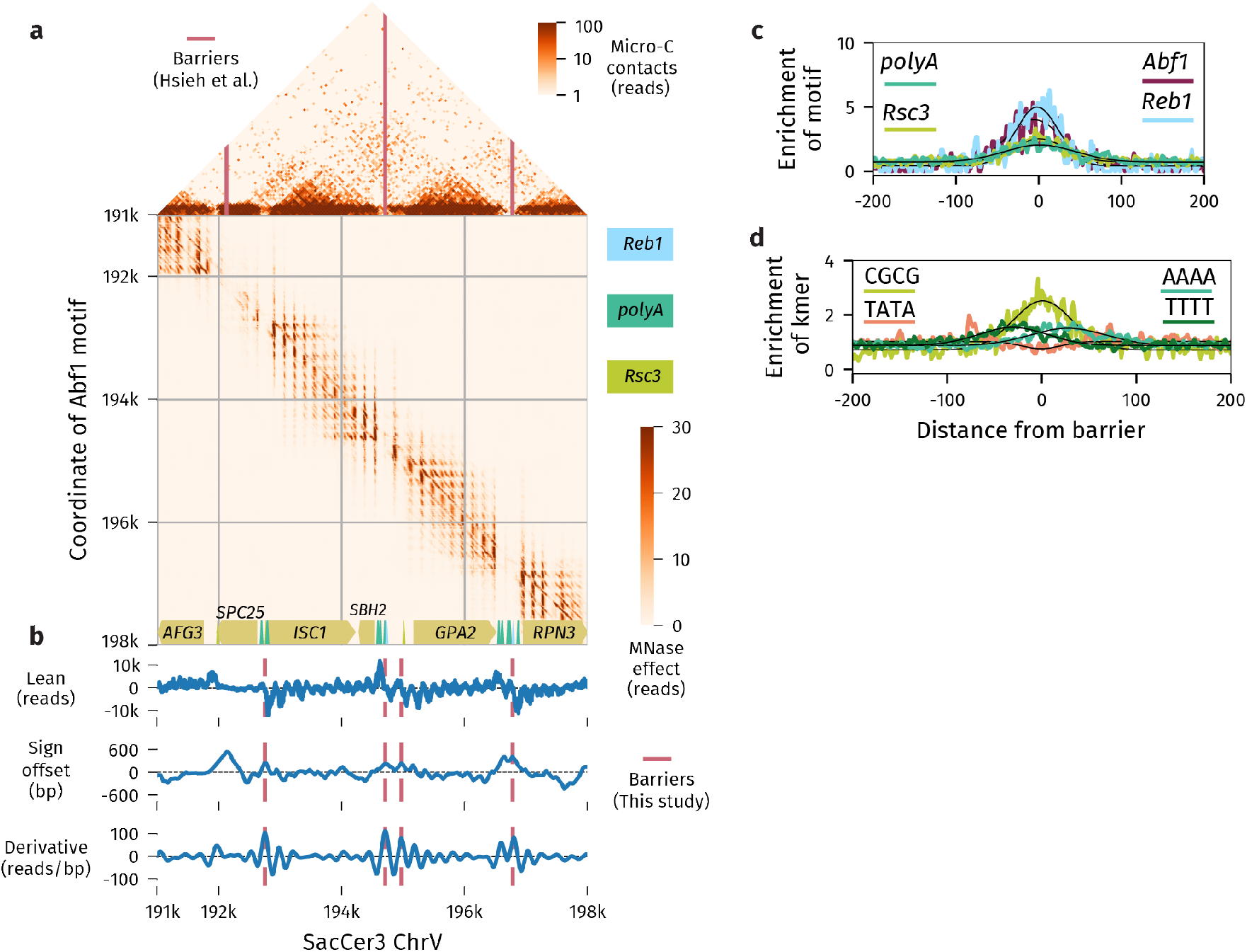
Additional barrier-like elements. (a) Effects of inserting an *Abf1* motif at the same region used in Oberbeckmann et al.[73]. Our results are concordant with the results in Figure 2 of their work. With the improved resolution afforded by our motif-scanning strategy, we can observe a single nucleosome between the promoters of *SBH2* and *GPA2* that seems to be an island unto itself. Further, our model shows that the NDR between *SPC25* and *AFG3* is not caused by an *Abf1* motif as suggested previously[73], but rather by an *Rsc3* motif at the end of the *SPC25* open reading frame. (b) Lean scores and called peaks for barrier elements in the same region. Pink lines indicate consensus peaks. The sign offset and derivative scores used to call barrier peaks are described in the methods. It is interesting to note that the derivative-based analysis fails to detect a barrier in the very wide NDR on the left of the region, but clearly shows the lonely nucleosome between *SBH2* and *GPA2* (c) Enrichment of four motifs at boundaries identified using Micro-C data[71]. The strong *Abf1* and *Reb1* motifs are highly enriched within a small window around the domain boundaries, while the *polyA* motif is more broad. (d) Enrichment of four kmers around boundaries identified previously[71]. The pattern is similar to that observed with our method, though our perturbation-based analysis is able to map these sequence elements at higher resolution.

**Table 1.** Performance comparison for OSKN model. We trained a BPReveal model using the same data as were used in Avsec *et al*[3]. We assess the quality of a model’s predictions against an experimental dataset using two metrics. The profile Jensen-Shannon divergence measures the similarity between the predicted and observed data at high resolution, while the Spearman correlation of the counts measures the performance of the model over the entire output window. Overall, BPReveal performs similarly to the previous state of the art in TF binding models.

|  |  | Counts Spearman correlation (higher is better) |  |  |  |  |  |
| --- | --- | --- | --- | --- | --- | --- | --- |
|  |  | Training set |  |  | Validation set |  |  |
| TF | strand | BPreveal | Avsec | $\Delta$ | BPreveal | Avsec | $\Delta$ |
| Oct4 | positive | 0.576 | 0.469 | **0.107** | 0.512 | 0.429 | **0.083** |
| Oct4 | negative | 0.577 | 0.464 | **0.113** | 0.513 | 0.425 | **0.088** |
| Sox2 | positive | 0.496 | 0.491 | **0.005** | 0.427 | 0.450 | -0.024 |
| Sox2 | negative | 0.500 | 0.488 | **0.012** | 0.424 | 0.443 | -0.019 |
| Klf4 | positive | 0.684 | 0.526 | **0.159** | 0.646 | 0.488 | **0.158** |
| Klf4 | negative | 0.684 | 0.529 | **0.154** | 0.647 | 0.491 | **0.156** |
| Nanog | positive | 0.616 | 0.661 | -0.045 | 0.579 | 0.642 | -0.064 |
| Nanog | negative | 0.618 | 0.665 | -0.047 | 0.580 | 0.648 | -0.068 |

|  |  | Profile Jensen-Shannon divergence (lower is better) |  |  |  |  |  |
| --- | --- | --- | --- | --- | --- | --- | --- |
|  |  | Training set |  |  | Validation set |  |  |
| TF | strand | BPreveal | Avsec | $\Delta$ | BPreveal | Avsec | $\Delta$ |
| Oct4 | positive | 0.696 | 0.680 | 0.017 | 0.696 | 0.679 | 0.018 |
| Oct4 | negative | 0.697 | 0.679 | 0.018 | 0.697 | 0.679 | 0.018 |
| Sox2 | positive | 0.758 | 0.753 | 0.005 | 0.757 | 0.751 | 0.006 |
| Sox2 | negative | 0.758 | 0.752 | 0.006 | 0.758 | 0.752 | 0.006 |
| Klf4 | positive | 0.675 | 0.642 | 0.033 | 0.677 | 0.644 | 0.033 |
| Klf4 | negative | 0.675 | 0.640 | 0.034 | 0.677 | 0.643 | 0.034 |
| Nanog | positive | 0.676 | 0.675 | 0.001 | 0.676 | 0.675 | 0.001 |
| Nanog | negative | 0.676 | 0.675 | 0.001 | 0.677 | 0.674 | 0.003 |

**Table 2.**
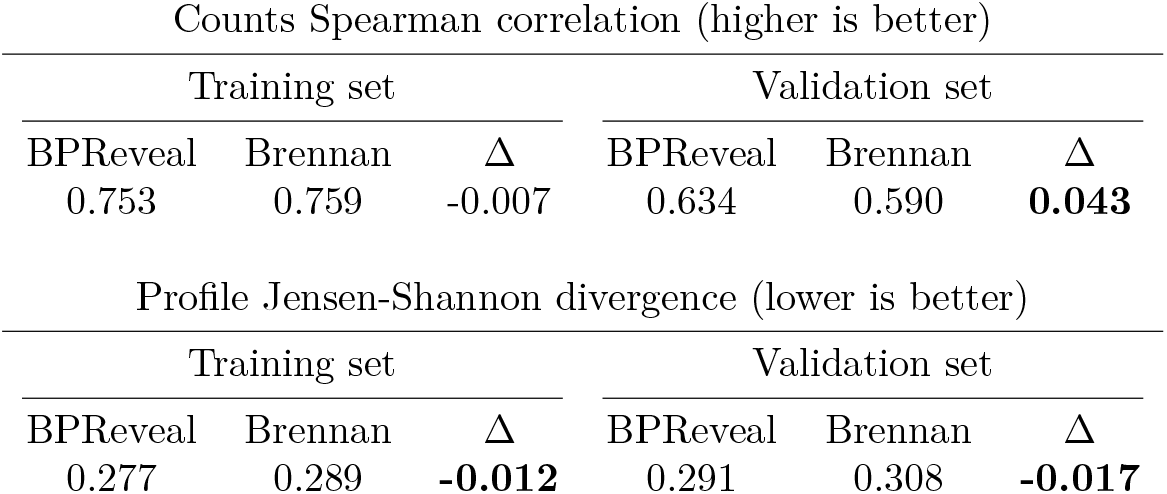
Performance comparison for ATAC-seq model. We trained a BPReveal model on the same data used in Brennan *et al*[15], and assessed the two models’ accuracy using the same metrics as in Extended Data Table 1. Our BPReveal model uses essentially the same architecture as ChromBPNet, and so the results are unsurprisingly very similar.

| Counts Spearman correlation (higher is better) |  |  |  |  |  |
| --- | --- | --- | --- | --- | --- |
| Training set |  |  | Validation set |  |  |
| BPreveal | Brennan | $\Delta$ | BPreveal | Brennan | $\Delta$ |
| 0.753 | 0.759 | -0.007 | 0.634 | 0.590 | **0.043** |
| Profile Jensen-Shannon divergence (lower is better) |  |  |  |  |  |
| Training set |  |  | Validation set |  |  |
| BPreveal | Brennan | $\Delta$ | BPreveal | Brennan | $\Delta$ |
| 0.277 | 0.289 | **-0.012** | 0.291 | 0.308 | **-0.017** |

**Table 3.** Performance data for BPReveal MNase model. These statistics are based on the 5^*′*^ endpoint predictions from the model trained on the data from Begley *et al*[48]. (Metrics for the 3^*′*^ endpoint predictions (not shown) are within 1% of the values in this table.)

| Metric | Training set | Validation set |
| --- | --- | --- |
| Profile Jensen-Shannon divergence (lower is better) | 0.235 | 0.260 |
| Profile Pearson (higher is better) | 0.856 | 0.815 |
| Counts Pearson (higher is better) | 0.717 | 0.626 |
| Counts Spearman (higher is better) | 0.711 | 0.612 |

**Table 4.**
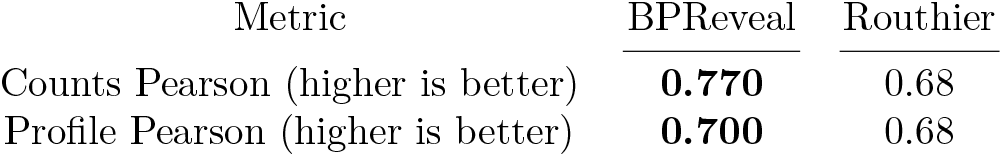
Performance comparison for Routhier MNase model. A BPReveal model was trained on the same data as the model in Routhier *et al*[25], and the BPReveal model performs as well or better than the previous state of the art. The correlation metric for the Routhier model is base-by-base for an entire region of a chromosome, and therefore there is no distinction between a profile and counts metric. Our metrics here are for the validation set.

| Metric | BPreveal | Routhier |
| --- | --- | --- |
| Counts Pearson (higher is better) | **0.770** | 0.68 |
| Profile Pearson (higher is better) | **0.700** | 0.68 |

